# Glioma-induced alterations in neuronal activity and neurovascular coupling during disease progression

**DOI:** 10.1101/763805

**Authors:** Mary Katherine Montgomery, Sharon H. Kim, Athanassios Dovas, Kripa Patel, Angeliki Mela, Nelson Humala, Hanzhi T. Zhao, David N. Thibodeaux, Mohammed Shaik, Ying Ma, Jack Grinband, Daniel S. Chow, Catherine Schevon, Elizabeth M. C. Hillman, Peter Canoll

**Author notes:** Co-corresponding authors Peter Canoll, Elizabeth Hillman. Equal contribution first authors (Mary Katherine Montgomery, Sharon H. Kim, Athanassios Dovas).

## Abstract

Diffusely infiltrating gliomas are known to cause alterations in cortical function, vascular disruption and seizures. These neurological complications present major clinical challenges, yet their underlying mechanisms and causal relationships to disease progression are poorly characterized. Here, we followed glioma progression in awake Thy1-GCaMP6f mice using in-vivo wide-field optical mapping to monitor alterations in both neuronal activity and functional hemodynamics. The bilateral synchrony of spontaneous neuronal activity in glioma-infiltrated cortex gradually decreased, while neurovascular coupling was also progressively disrupted compared to uninvolved cortex. Over time, mice developed diverse patterns of high amplitude discharges and eventually generalized seizures that begin at the infiltrative margin of the tumors. Interictal and seizure events exhibited positive neurovascular coupling in uninfiltrated cortex, however glioma-infiltrated regions exhibited inverted hemodynamic responses driving seizure-evoked hypoxia. These results reveal a landscape of complex physiological interactions occurring during glioma progression and present new opportunities for exploring new biomarkers and therapeutic targets.

**Highlights:**

- Glioma disrupts neural synchrony between bilateral cortical regions.
- WFOM reveals frequent interictal discharges and seizures during glioma progression.
- Tumor development is accompanied by local changes in neurovascular coupling.
- Altered neurovascular coupling drives hypoperfusion of the tumor during seizures.

## Introduction

Gliomas are primary brain neoplasms characterized by diffuse infiltration of glioma cells into surrounding brain tissue, where they interact with neurons, astrocytes and blood vessels (Cuddapah et al., 2014; Torres and Canoll, 2019). Glioma patients often present with neurological symptoms that result from functional alterations in the infiltrated cortex. Seizures, the most common neurological symptom, afflict >80% of low-grade glioma patients and 50-60% of high-grade glioma patients and can contribute significantly to the deterioration of cognitive function (Klein et al., 2003; Ruda et al., 2010). Recent studies have further suggested that increased seizure activity may contribute to increased tumor growth and progression (Johung and Monje, 2017) (Vecht et al., 2014) (Robert et al., 2015). The clinical management of glioma-associated seizures is confounded by complex interactions between standard epilepsy drugs, therapies targeting tumor progression and tumor pathophysiology itself (de Groot et al., 2012; van Breemen et al., 2007). Currently, the most effective therapeutic approach to ameliorating seizures in glioma patients is by maximizing tumor resection to include the infiltrating tumor margins (Pallud et al., 2014). However, when glioma patients fail therapy and their tumors recur, they usually present with new or worsening seizures (Smits and Duffau, 2011). These clinical observations suggest that glioma-induced epileptogenesis is a dynamic process caused by the infiltration of glioma cells into the cortex, which leads to progressive alterations in focal cortical activity. Patients with seizures also develop brief interictal epileptiform discharges that do not cause overt behavioral manifestation, but can occur much more frequently than observable seizures (Selvitelli et al., 2010). However, the causes and cognitive effects of these smaller interictal events (IIEs) remain controversial (Avanzini et al., 2013; Faught et al., 2018), and their relationship to tumor progression is not well characterized. The complex interplay between glioma progression and aberrant neural activity is poorly understood, and yet underlies many aspects of tumor progression, directly affecting the clinical care of almost all low-grade glioma patients.

Glioma cells have also been observed to interact closely with blood vessels, migrating along them as conduits during tumor spread, and disrupting their structural and functional integrity (Farin et al., 2006) (Watkins et al., 2014). These vascular interactions prompt the question of whether tumors could impact the coupling between neuronal activity and vascular dynamics during glioma-infiltration. Such neurovascular interactions could contribute to the disturbed functional and metabolic state of the brain during tumor progression, particularly in the context of abnormal neural events such as IIEs and seizures. Although not studied in the setting of gliomas, epileptic seizures have been shown to have profound hemodynamic effects that results in focal and transient increases in cerebral perfusion (Ma et al., 2013; Schwartz, 2007; Zhao et al., 2009). Glioma-related changes in neurovascular coupling could also affect the hemodynamic signals measured using functional magnetic resonance imaging (fMRI). Understanding these effects is especially important since the primary clinical use of fMRI in the setting of glioma is pre-surgical planning for tumor resections, seeking to identify the location of key functional brain regions to be preserved or removed (Genetti et al., 2013; Petrella et al., 2006). Tumor-dependent alterations in neurovascular coupling could lead to misinterpretation of fMRI data, and errors in the localization of functional boundaries. Several recent human resting–state fMRI studies have also shown that gliomas can cause focal changes in the blood oxygen level dependent (BOLD) signals in the tumor and surrounding infiltrated cortex, characterized by a loss of synchrony between the tumor and the contralateral hemisphere (Bowden et al., 2018) (Chow et al., 2016) (Agarwal et al., 2016). Although, these studies could not disambiguate whether changes in BOLD were caused by altered neuronal activity, neurovascular coupling or both, they suggest that vascular signals could be helpful biomarkers of tumor progression.

To understand these diverse, dynamic physiological interactions between tumor progression, neuronal activity and hemodynamics, we need to monitor the in-vivo functioning brain and assess the alterations of both neuronal activity and neurovascular responses during glioma progression. Here we combined novel methods for in-vivo, longitudinal wide-field optical mapping (WFOM) in awake behaving Thy1-GCaMP6f mice (Ma et al., 2016a) with an orthotopic model of glioma, characterized by diffuse infiltration of the cerebral cortex with perivascular patterns of invasion, and intermingling of tumor cells with neurons. Neuronal activity and hemoglobin oxygenation dynamics were simultaneously imaged across both hemispheres of the dorsal cortex throughout the course of disease progression within individual mice. This imaging approach enabled direct observations of longitudinal changes in the properties of spontaneous and evoked neuronal events across the cortex including intracortical desynchronization, progressively worsening IIEs events and eventually spontaneous seizures. These data also afforded the ability to explore the way in which these different neuronal events were coupled to changes in hemodynamics as a function of disease progression. Our analysis revealed significant neurovascular disruption within tumor-burdened regions, a finding that has important implications for interpretation of fMRI data acquired for resection guidance, while also suggesting ways in which altered coupling could contribute to exacerbation of tumor-related damage and even hasten tumor progression. These results provide new insights into the progressive interplay between glioma invasion and cortical function, while demonstrating that WFOM is a valuable tool for characterizing disease and potentially screening for new therapeutic targets to reduce the morbidity and mortality of glioma.

## Results

Wide field optical mapping (WFOM) was used to longitudinally image GCaMP6f fluorescence and hemodynamics across the cortex of three adult Thy1-GCaMP6f mice following intracerebral injection of PDGFA^+^/ TP53^-/-^ mouse glioma cells into the subcortical white matter of right frontal cortex. Thy1-GCaMP6f mice exhibit widespread changes in cortical fluorescence corresponding to changes in intracellular calcium levels within excitatory neurons of layers 2/3 and 5. A thinned-skull cranial window was placed at the same time as tumor initiation, and after recovery, the mice were imaged during sessions lasting up to 2 hours in which they were head-restrained but awake, starting at 4 days post injection (DPI) and continuing up to 36 DPI. WFOM images were acquired using a camera focused onto the cranial window with a sequence of LEDs illuminating to record green fluorescence from GCaMP6f under blue illumination, and then green and red reflectance to enable calculation of changes in oxy-, deoxy- and total hemoglobin (HbO, HbR and HbT=HbO+HbR) as detailed in STAR methods. Reflectance measurements were also used to correct GCaMP6f fluorescence measurements for the effects of changes in hemoglobin absorption (Ma et al., 2016a). All animals expired at the end of the study and their brains were extracted for histological processing. Additional mice were implanted with tumors in the same way to permit more comprehensive histological analysis at interim time-points.

### Diffuse infiltration of glioma into the cortex of Thy1-GCaMP6f mice leads to progressive alterations in neurons and blood vessels

Histological analysis demonstrated that gliomas produced using this method showed a progressive and diffuse infiltration into the cortex, recapitulating the patterns of infiltration seen in human gliomas (Figure 1A). Perineuronal satellitosis was prominent, particularly at the infiltrative margins of the tumor (Figure 1B). The density of neurons was significantly decreased in the highly cellular core of the tumors, and to a lesser extent at the infiltrative margins (Supplementary Figure S1A-B). Infiltrating glioma cells also accumulated near blood vessels (stained using the isolectin IB4; Figure 1C).

**Figure 1.**
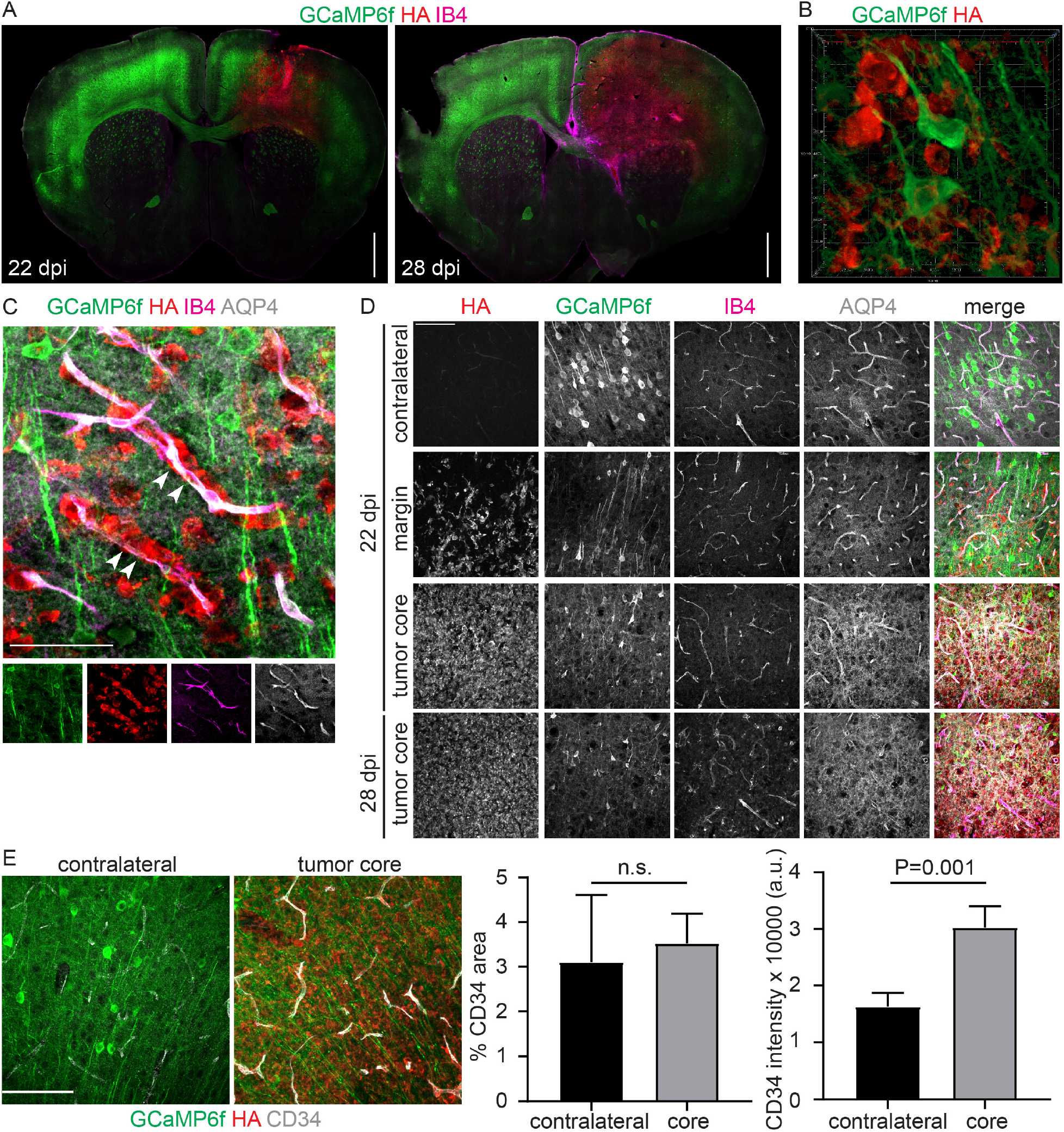
Immunohistochemical analysis reveals diffuse infiltration of the cortex and progressive disruption of the vasculature within the highly cellular core of the tumor. (A) Coronal sections obtained from glioma-bearing mice at 22 and 28 days post injection (DPI) shows the distribution of glioma cells (HA; red), GCaMP6f^+^ neurons (green) and IB4-labeled vessels (magenta). Stitched images show a single confocal plane. Bar, 1000 µm. B) 3D rendering of GCaMP6f^+^ neurons (green) surrounded by infiltrating glioma cells (red). Image dimensions: 80 µm, 80 µm, 15 µm (X,Y,Z). C) Infiltrating glioma cells (red) migrate along blood vessels (arrowheads), with AQP4 (white) remaining closely associated with IB4+ vessels (magenta). Maximum intensity projections of confocal stacks. Bar, 50 µm. D) Representative fields of glioma at the indicated time points and locations. AQP4 association with IB4^+^ vessels is intact in the contralateral (uninfiltrated) cortex and at the infiltrative margins of the tumor, but is disrupted in the tumor core. Bar, 100 µm. E) Contralateral cortex and tumor core of a 28 DPI tumor stained with CD34. Maximum intensity projections of confocal stack. Bar, 100 µm. Graphs (mean + standard deviation) show that the density of CD34+ vessels in the tumor is not significantly altered, however the intensity of CD34 immunoreactivity is significantly higher in the tumor vessels compared to contralateral cortex. Two-tailed, unpaired t-test was used to calculate significance.

Notably, immunofluorescence analysis for the water channel aquaporin 4 (AQP4), which in healthy brain tissue is discretely localized in astrocyte end-feet contacting vessels (Mader and Brimberg, 2019), revealed that astrocyte end-feet remained closely associated with vessels at the infiltrative margins (Figure 1D). In contrast, in the highly cellular core of the tumor, AQP4 became diffuse and disorganized and was no longer tightly associated with vessels (Figure 1D). Since IB4 also recognizes phagocytes, we examined specific changes in the vasculature of the glioma-infiltrated cortex using CD34, a glycoprotein present on vascular endothelial cells (Nielsen and McNagny, 2008). Confocal images acquired in the highly cellular core of the tumor showed that vessels had an abnormal morphology and were encased by glioma cells (Figure 1C), and while there was no significant change in the overall density of CD34+ vessels, the levels of CD34 immunoreactivity in the vasculature within the tumor core was significantly increased (Figure 1E). Similar changes in AQP4 and CD34 are seen in surgical samples of human gliomas (Noell et al., 2012) (Smith and Verkman, 2015) (Clara et al., 2014), and have been proposed to result from tissue hypoxia (Nigim et al., 2015) (Mou et al., 2010). Two-photon images acquired post-mortem from the infiltrating margins of the tumors from the mice that underwent in vivo imaging allowed us to interrogate the patterns of glioma infiltration through layers 2/3 and 5. In these mice, glioma cells intermingled with GCaMP6f^+^ neurons (Supplementary Figure S1C and Supplementary Video S1), and appeared to migrate along the abluminal surface of blood vessels in close association with AQP4+ astrocyte endfeet (Supplementary Figure S1D). These findings reveal progressive disruption of neuronal and vascular organization in the glioma-bearing cortex, which is most severe in the highly cellular tumor core, but remains relatively intact at the infiltrated margins of the tumor.

### Resting state neuronal and hemodynamic activity becomes less bilaterally correlated in tumor regions during glioma progression

Figure 2 shows a sequence of longitudinal in-vivo WFOM recordings in one animal during the progression of tumor growth in the anterior right side of the cortex. From baseline GCaMP6f fluorescence measurements, a gradually expanding dark area is visible around the tumor cell injection site in the right anterior quadrant. Corresponding green light reflectance images do not show the same darkening, confirming that this effect corresponds to decreasing GCaMP6f fluorescence and not increasing absorption, for example from increased vascular density (Figure 2A). This GCaMP6f signal decrease is consistent with the progressive loss of neurons, which is most pronounced in the highly cellular core of the tumor (as shown in Supplementary Figure S1).

**Figure 2.**
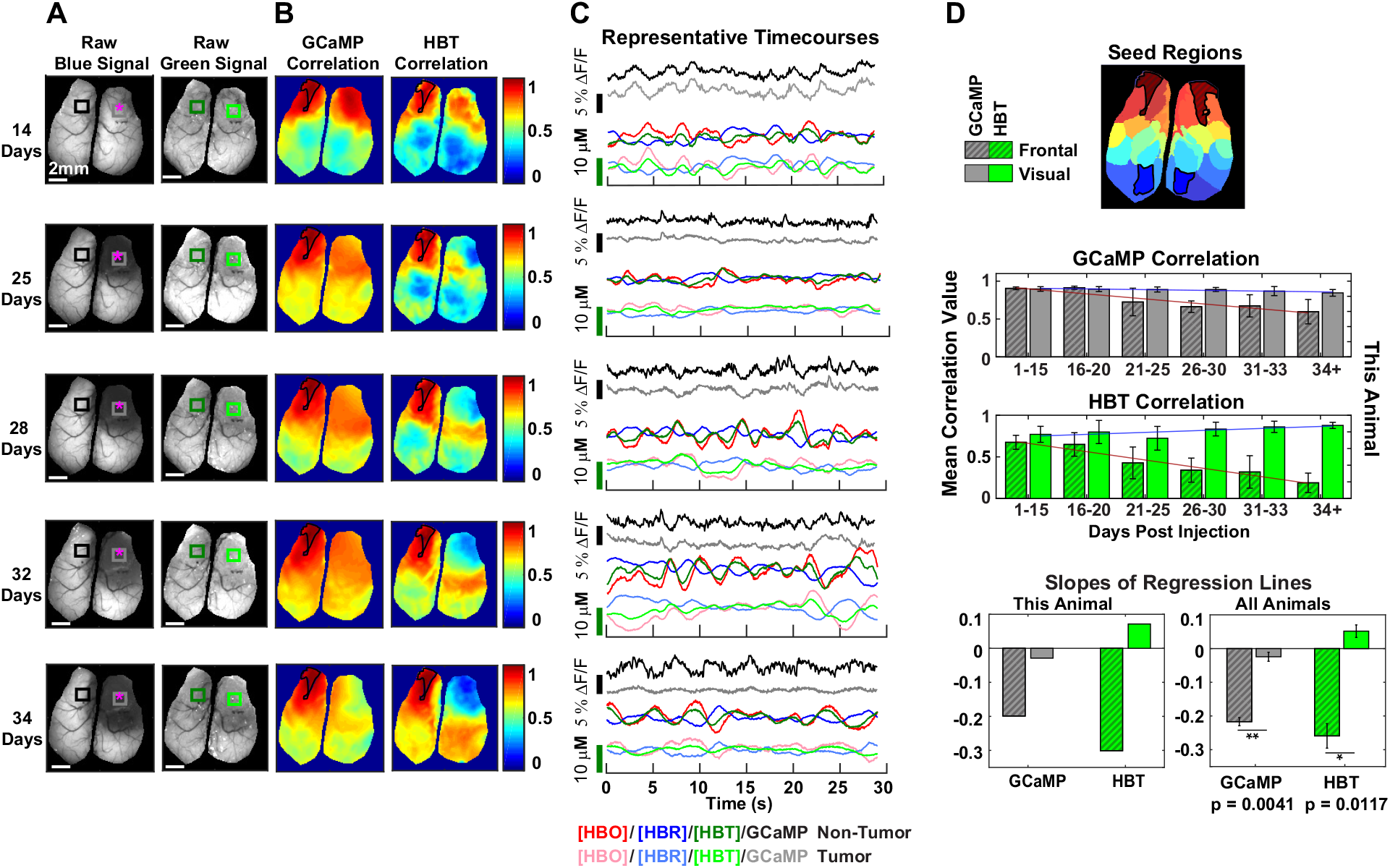
Correlation of GCaMP6f and hemodynamic signals between tumor and non-tumor regions during tumor progression (Mouse 2). (A) Images of raw fluorescence signal showing progressive loss of GCaMP6f fluorescence with no corresponding loss of hemodynamic signal. Magenta asterisks indicate approximate site of glioma injection. (B) Maps of correlation to a seed region, outlined in black, for GCaMP6f and hemodynamic data. Note that there is high HBT correlation at the tumor margin, even at late timepoints, in contrast to the decreased correlation evident in the tumor core. (C) Individual time courses taken from representative runs and averaged across regions in the tumor and non-tumor areas of the brain. (D) Top: Map of k-means seed regions used in correlation analysis with frontal cortex (dark red diagonal hatch line fill) and visual cortex (dark blue). Center: Bar plots of both GCaMP6f and hemodynamic correlation in frontal (gray with diagonal hatch line fill) and visual regions (gray fill) as tumor progresses, shown for one representative animal. Bottom: Slopes of regression lines for GCaMP6f and hemodynamic correlation values shown for one representative animal and across all animals (error bars are standard error across animals). Significance across all animals was calculated using paired, two-sample, two-tailed t-tests between non-tumor and tumor regions for each channel at significance of p < 0.05 (*) and p < 0.005 (**).

Figure 2C shows example time series from the tumor and the contralateral anterior region. At early time points, neuronal activity on both sides of the cortex are clearly well-correlated, and the same is true for bilateral cortical hemodynamics. However, as the tumor progresses, neuronal activity in the tumor region can be seen to decrease both in terms of its % amplitude, and its high frequency content compared to the contralateral region (gray and black traces). Hemodynamic activity in the tumor region also shows a progressive attenuation of fluctuations, particularly in HbT. This effect can also be seen in the representative real-time WFOM data movies (Supplementary Video S2).

To assess whether these changes correspond to decorrelations of activity, Pixel-by pixel Pearson’s correlation analysis was performed to map the level of synchrony between cortical regions. The maps shown in Figure 2B depict the correlation of signals relative to the anterior left frontal seed region (contralateral to the tumor) for both GCaMP6f and HbT. These correlation maps show a gradual decrease in regional correlation of both neuronal and hemodynamic signals at the core of the tumor as it progresses, with more significant changes visible in the HbT correlation maps, a pattern that was consistent across all animals (Supplementary Figure S2). Notably, the region corresponding to the margin of the tumor shows a higher HBT correlation with the contralateral cortex than the tumor core, even though both regions show neuronal loss evident from the low GCaMP signal (Figure 2A and 2B). These findings suggest that the infiltrating margin of the tumor is functionally distinct from the tumor core, particularly with regards to hemodynamic activity.

To quantify these trends, bilateral correlations between 4 brain regions were compared: the anterior right frontal cortex (tumor region), the anterior left frontal cortex (contralateral to the tumor region), and visual cortex areas ipsilateral and contralateral to the tumor (Figure 2D). While almost no changes in bilateral correlation were seen for anterior (non-tumor-bearing) regions, significant and systematic decreases in bilateral correlation were seen for the frontal regions as the tumor progressed unilaterally. The difference in the slope of this correlation trend between tumor and non-tumor regions was significant, and more pronounced for hemodynamics than for GCaMP6f signals (Figure 2D). A consistent effect was seen in all 3 animals (see Supplementary Figure S2). However, we note that GCaMP6f analysis is impacted by the gradual decrease in signal to noise caused by the significant loss of GCaMP fluorescence in the tumor as it progresses.

The significance of these decorrelations relates to the field of resting state fMRI. The BOLD signals detected in resting state fMRI correspond to seemingly random fluctuations in [HbR], which have been found to be correlated within brain-wide functional connectivity networks (Raichle et al., 2001). Recent studies have demonstrated that brain-wide patterns of neuronal activity are predictive of hemodynamic fluctuations, suggesting that resting state functional connectivity depicts an important property of brain-wide neuronal dynamics (Ma et al., 2016b). Our results demonstrate that glioma infiltration disrupts the otherwise strong bilateral synchrony of both resting-state neuronal and hemodynamic fluctuations, a feature consistent with a recent study that noted the ability of resting state fMRI analysis to delineate glioma tumor margins (Chow et al., 2016).

### Tumor growth systematically alters local resting state neurovascular coupling

The results above demonstrate that the growing tumor disrupts the synchrony of patterns of spontaneous neuronal activity and hemodynamics in the awake mouse cortex (Ma et al., 2016b). However, although this effect appears to be stronger for hemodynamics than neuronal activity, this analysis does not directly address the question of whether the coupling between neuronal activity and hemodynamics within each region is also altered by tumor burden.

It is well-described that in the normal brain, stimulus-evoked neuronal activity generates a stereotyped increase in local blood flow, driving an increase in HbT and HbO and a decrease in HbR. The positive BOLD response to a stimulus detected in fMRI corresponds to this decrease in [HbR], providing a surrogate measure of neural activity (Hillman, 2014; Kwong et al., 1992). To explore the effect of glioma progression on the cortical response to external stimulation, in every WFOM imaging session, mice underwent a sequence of 60 trials in which they received 5 second tactile whisker stimulation to either their right or left whisker pad at 30 second intervals. As shown in Supplementary Figure S3 averaged cortical responses (excluding trials in which the mice ran) did not reveal significant differences in either the cortical neuronal response to the whisker stimulus, nor the corresponding localized hemodynamic response. However, as shown in Supplementary Figure S3 A-B, the whisker barrel cortex is located posterior to the tumor region, and is outside the infiltrating margin of the tumor, even at the latest time-points measured, and thus this stimulus did not directly probe an evoked response within the tumor-invaded cortex. Nevertheless, this result suggests that the presence or progression of the tumor does not have a strong global effect on stimulus evoked-neurovascular coupling in non-tumor regions.

Even without a stimulus-evoked response within the tumor region, however, it is still possible to quantify the coupling between hemodynamic and neuronal activity within the tumor region if we rely on analysis of the relationship between spontaneous neural and hemodynamic events. To do this, we performed deconvolution analysis, as previously described (Ma et al., 2016b), to estimate the coupling relationship between spontaneous fluctuations in neuronal and hemodynamic signals for different regions of interest across the cortex as the tumor progressed. Analysis was performed using only epochs of WFOM data in which the animal did not noticeably move (based on simultaneous behavioral monitoring; see STAR methods for full details of methods including hemodynamic cross-talk correction). Analysis was performed using signals extracted from regions of interest spanning both sides of the cortex, from 14 DPI onwards in all mice. Deconvolution was used to estimate HbO, HbR and HbT hemodynamic response functions corresponding to the system’s impulse response to a neuronal event.

In posterior brain regions, further from the tumor center, normal stereotyped hemodynamic response functions were observed, exhibiting an expected increase in HbT and HbO with a decrease in HbR consistent with stimulus-evoked functional hyperemia and earlier results in normal mice (Ma et al., 2016b). However, in ROIs in tumor regions, a progressive pattern of changing hemodynamic response functions were observed. Initially, the response became attenuated and delayed, and dominated by an initial condition in which HbR levels were high. The deconvolution model was adjusted based on this observation to introduce a time-shift between the neuronal and hemodynamic data to determine whether any coupled hemodynamic changes occur before the peak of neuronal events. This analysis indeed revealed pre-neural event patterns of hemodynamics varying from apparently pre-emptive decreases in HbT, to later periods in which hyperemia appears to precede the neuronal response (Figure 3). It should be noted that the decreasing amplitude of the GCaMP6f fluorescence signal within the tumor led to increased noise at later time-points, potential cross-talk confounds and thus lower confidence in the precise temporal shapes of hemodynamic response functions within the tumor itself. In some epochs, clear hemodynamic oscillations were also observed. These effects are evident in the convolution correlation metrics in Figure 3C, which show the correlation between the original red and green reflectance data, and the model-fit corresponding to correlation between the HRF and neural signal. Lower correlation values at later time-points in tumor-infiltrated regions suggest that a simple linear HRF model is less accurate within the tumor. However, the unusual HRF patterns found were consistent across days, gradually spreading to different cortical regions as the tumor progressed and invaded larger regions of cortex, and were generally reproduced in all mice, as shown in Figure 3D and Supplementary Figure S4.

**Figure 3.**
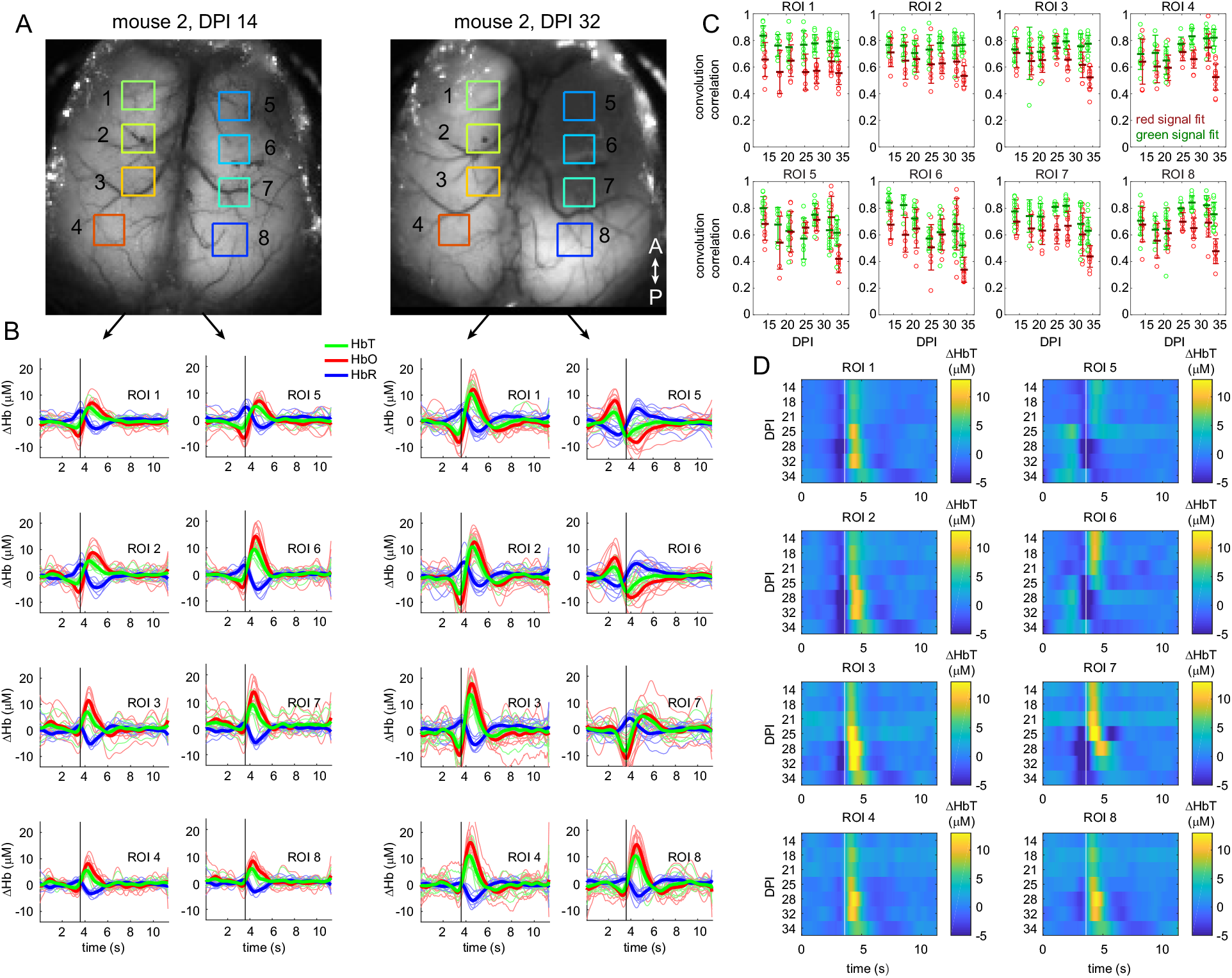
Analysis of regional changes in neurovascular coupling during tumor progression by deconvolution of resting state neural activity from hemodynamics. (A) shows the brain of mouse 2 at early and late time-points with regions of interest spanning the cortex contralateral and ipsilateral to the tumor. (B) shows hemodynamic response functions (HRFs) calculated using deconvolution of resting state epochs (see STAR methods), with fine lines showing individual trial results, and bold lines showing the average for each imaging session. The left two columns show ROIS 1-8 at 14 days past injection (DPI) showing bilaterally consistent responses in which HbT increases following the black line which denotes the peak of a spontaneous neural event. The second two columns show HRFs from the same 8 ROIs at 32 DPI. Here, the contralateral side has consistent hemodynamic responses, whereas the tumor-affects side shows profound changes including hyperemia prior to the event followed by an evoked decrease in HbT within the tumor, and an intermediate response at the tumor boundary where the neural event occurs during a distinct decrease in HbT. (C) shows correlation coefficients of each trial between the original red and green signals, and signals predicted by the convolution model Later days for tumor regions show degraded correlations suggesting worsening fits to a linear HRF fit. Lower correlation values for red compared to green reflectance correspond to worse noise on raw red signals. (D) depicts aggregate average HRF results for all measurement sessions over the same 8 ROIs. The attenuation of the hemodynamic response within the gradually progressing tumor region is clear, along with the trends of initial low HbT and eventually pre-event hyperemia. The same analysis is shown for mice 1 and 3 in Supplementary Figure S4.

Our interpretation of these observations is that stimulus-evoked neuronal activity, such as a whisker flick, cannot evoke an anticipatory change in hemodynamics, and thus the classic stimulus-evoked HRF must always start from a baseline state. However, the activity spontaneously generated by the brain in periods of rest is not well characterized, and may itself be abnormal in the tumor and surrounding regions. If neural events are caused by local competing effects, such as excitation and inhibition, there could readily be GCaMP6f-independent neural or even non-neuronal metabolic effects driving coupled hemodynamics prior to the occurrence of a large GCaMP6f+ increase in neuronal activity.

A further feature revealed by this analysis was a slight, but noticeable change in the HRF throughout the brain at the latest stages of disease. Here, the HRF became noticeably lower in amplitude and smoothed out in time. These results provide strong evidence that the presence and growth of the tumor has a significant effect on coupling between neuronal activity and hemodynamics.

### Interictal events accompany tumor progression and evoke altered neurovascular coupling in regions of tumor infiltration

Over the ∼35 days of longitudinal imaging, the glioma-bearing mice developed increasingly abnormal neuronal activity, in particular, spontaneous high-amplitude discharges seen as sharp, high amplitude increases in GCaMP6f fluorescence with associated hemodynamic changes. We interpret these patterns to be tumor-related interictal events (IIEs) (Figure 4B-C and Supplementary Video S3). Events sometimes occurred in quick successive trains, while some occurred individually. Some events extended across all regions of the cortex, while others appeared to remain more local (Figure 4B-C). Spatially-dependent onset time analysis of successive IIE events within a single trial at DPI 35 shows widely varying initial locations, patterns and speed of spread, although involvement of the tumor margin is clear in many cases (Figure 4D). The occurrence of these events across all mice and imaging sessions is depicted in detail in Supplementary Figure S5 and summarized in Figure 4E. The overall frequency of IIEs correlated with the gradual growth of the tumor (with IIEs starting after 14 DPI). These events are not seen in WFOM recordings of healthy Thy1-GCaMP6f mice (Ma et al., 2016b).

**Figure 4.**
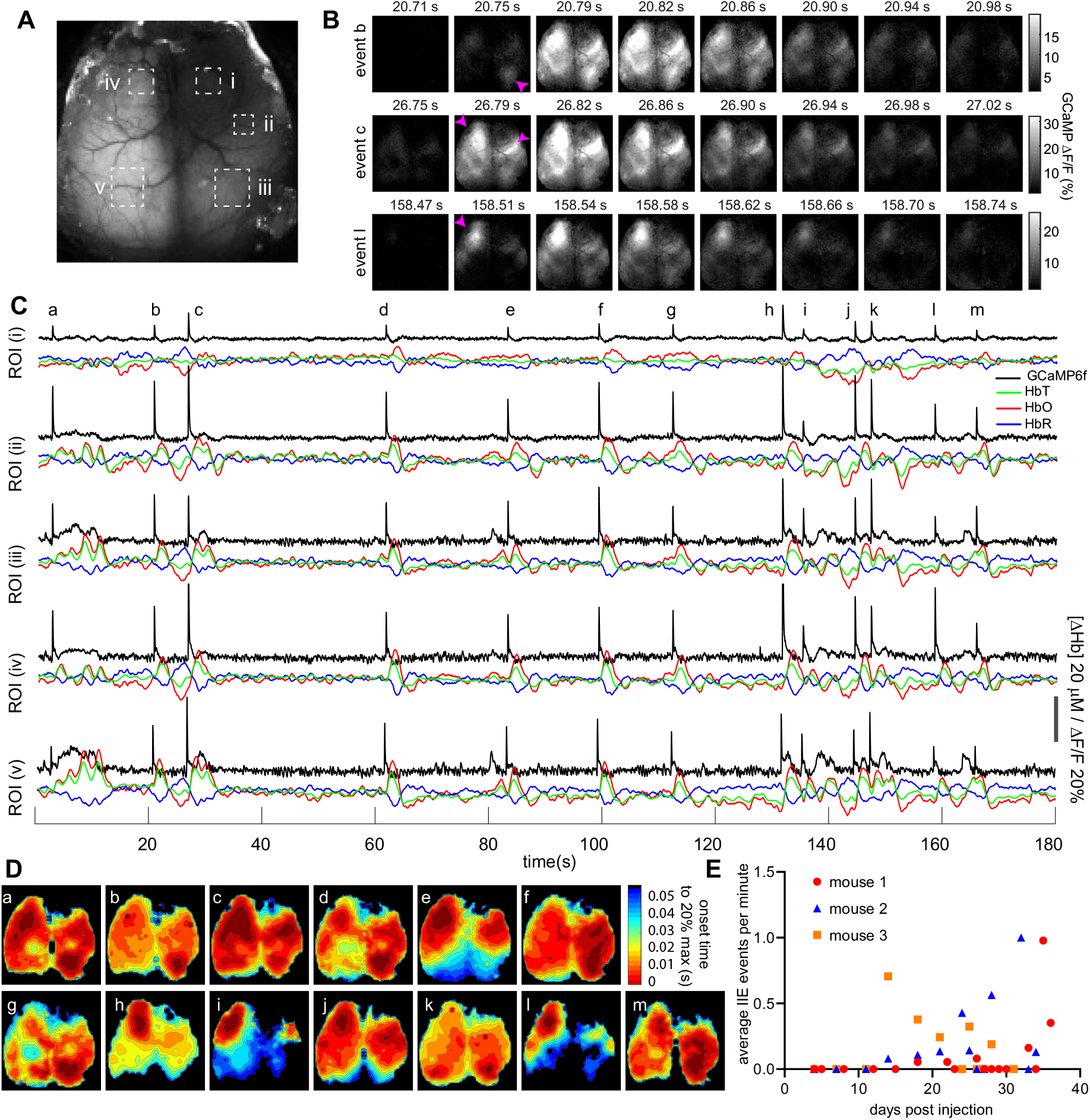
Spatial variation of initiation and propagation of interictal events (Mouse 1) (A) Raw fluorescence and (B) ΔF/F image time-series of 3 epochs from a 180 s resting state trial recording containing 13 spatially distinct interictal events (collected 35 days post injection). Magenta arrows indicate areas of interictal event initiation. (C). Time-courses of GCaMP6f fluorescence, HbO, HbR and HbT extracted from the 5 regions of interest (ROIs) in A corresponding to i-v the tumor, tumor margin, posterior region on the tumor side, anterior non-tumor and posterior non-tumor regions respectively. The lower amplitude and altered hemodynamic responses in the tumor region are clearly visible, as well as differing relative amplitudes of the IIEs in each region for each event, reflecting local or global spread. (D) Contour plots showing the onset time to 20% of the maximum for each of the 13 events (a-m) denoted in (C). Red regions correspond to t=0 initiation locations (see STAR methods). (E). Frequency of events for each mouse (all identified events divided by the total time of resting state imaging per session) over the course of all imaging sessions for all mice. Sessions including seizure events occur on DPI 34 (mouse 2) and 35 (mouse 1) are indicated with an asterisk. There is a significant increase in relative frequency of events over time, (R=0.3282, P-Value =0.036177).

The presence of spontaneous IIEs afforded the opportunity to assess how vascular responses are coupled to IIEs, both in tumor-burdened regions and the rest of the cortex. Analysis was performed using spike-triggered averaging centered on the peak of identified IIEs and results are shown for tumor-infiltrated versus distant cortex in Figure 5 and Supplementary Figure S6. In regions distant from the tumor, hemodynamic coupling to IIEs was found to be positive, corresponding to an increase in perfusion leading to increased HbT and HbO and a decrease in HbR, similarly to a ‘positive BOLD’ hemodynamic response to stimulus (compare to Supplementary Figure S3). However, similarly to the deconvolution results above, which excluded periods with notable IIEs, in glioma-infiltrated cortex, the same IIEs revealed a very different hemodynamic response whose peak is significantly delayed and attenuated compared to healthy cortex, and includes a slight inverted hemodynamic response which begins before the peak of the IIE. These patterns were found to be consistent across individual events, in individual animals and between animals. (Figure 5B-D, and Supplementary Figure S6B-D). These delays and reduced positive amplitude of hemodynamic responses to IIEs in tumor regions (relative to the amplitude of local GCaMP6f signals) suggests that this neurovascular impairment is likely under-serving the metabolic needs of the tumor region during these high amplitude neuronal events.

**Figure 5.**
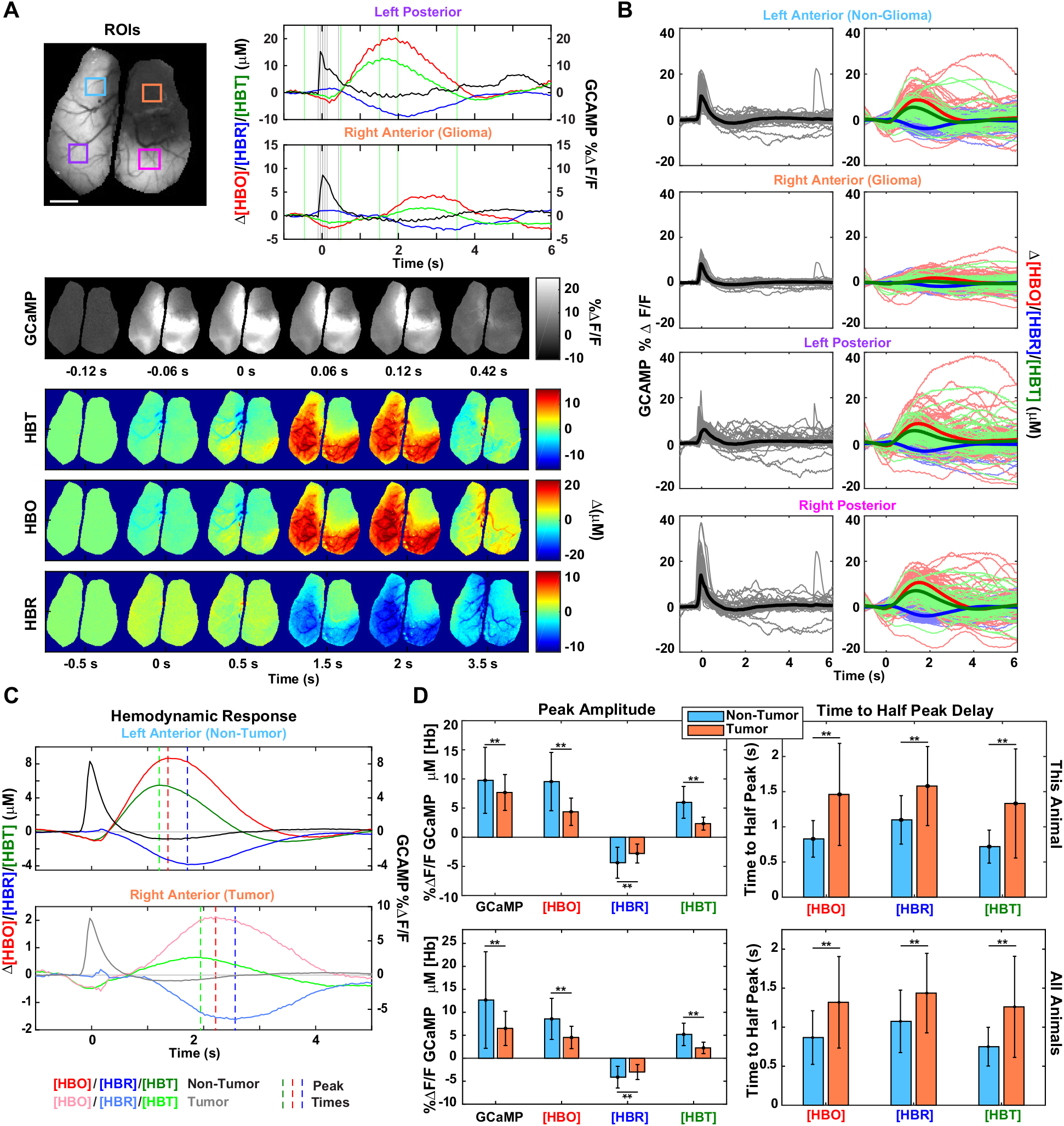
Region-specific alterations in neurovascular coupling during interictal events (Mouse 2). (A) Top Left: Raw fluorescence image marked with relevant regions of interest. Top Right: Time series taken from a representative interictal event (34 days post injection) and averaged across pixels in the left posterior and right anterior (tumor) regions. Relevant frames of interest for GCaMP6f marked with gray vertical lines and for hemodynamics with green vertical lines. Bottom: Spatial patterns of GCaMP6f and hemodynamic data shown for relevant frames of interest during the event. (B) Temporally aligned time series for all interictal events across all sessions (all days) in one representative animal averaged across pixels in various regions of interest and shown for both GCaMP6f and hemodynamic data, with the average plotted in bold. (C) Temporally aligned time series, averaged across all events across all sessions (all days) in same representative animal, showing the hemodynamic responses in non-tumor and tumor regions, averaged across trials and shown with vertical bars at peak times. The vertical axes for the GCaMP6f and hemodynamic right anterior (tumor region) time courses are scaled differently due to amplitude differences. (D) Bar plots of peak amplitude of GCaMP6f and hemodynamic data (left column) and time to half peak of hemodynamic data (right column), shown for one representative animal (top row) and across all animals (bottom row). Error bars are standard deviation across trials for one representative animal, and standard deviation across all animals. Significance across all animals was calculated using paired, two-sample, two-tailed t-tests between non-tumor and tumor regions for each channel at significance of p < 0.05 (*) and p < 0.005 (**).

### Spontaneous seizures in glioma-bearing mice reveal differences in the neurovascular response in tumor versus distant cortical regions

In addition to IIEs, our WFOM imaging sessions captured several spontaneous generalized seizure events in the late stage of tumor progression in two of the mice as shown in Figures 6 and 7 and Supplementary Videos S4 and S5. Our real-time recordings of the cortical representations of these spontaneous seizure events permitted observation of the progression and spread of high amplitude neuronal activity and corresponding vascular responses, as well as concomitant hemodynamic responses to the seizure activity. In all seizures observed, the event began as a series of high amplitude spikes consistent with interictal event characteristics, with highest signal intensities initially seen at the infiltrative margins of the tumor. This high intensity neuronal activity can then be seen to spread to the surrounding cortex in both hemispheres.

**Figure 6.**
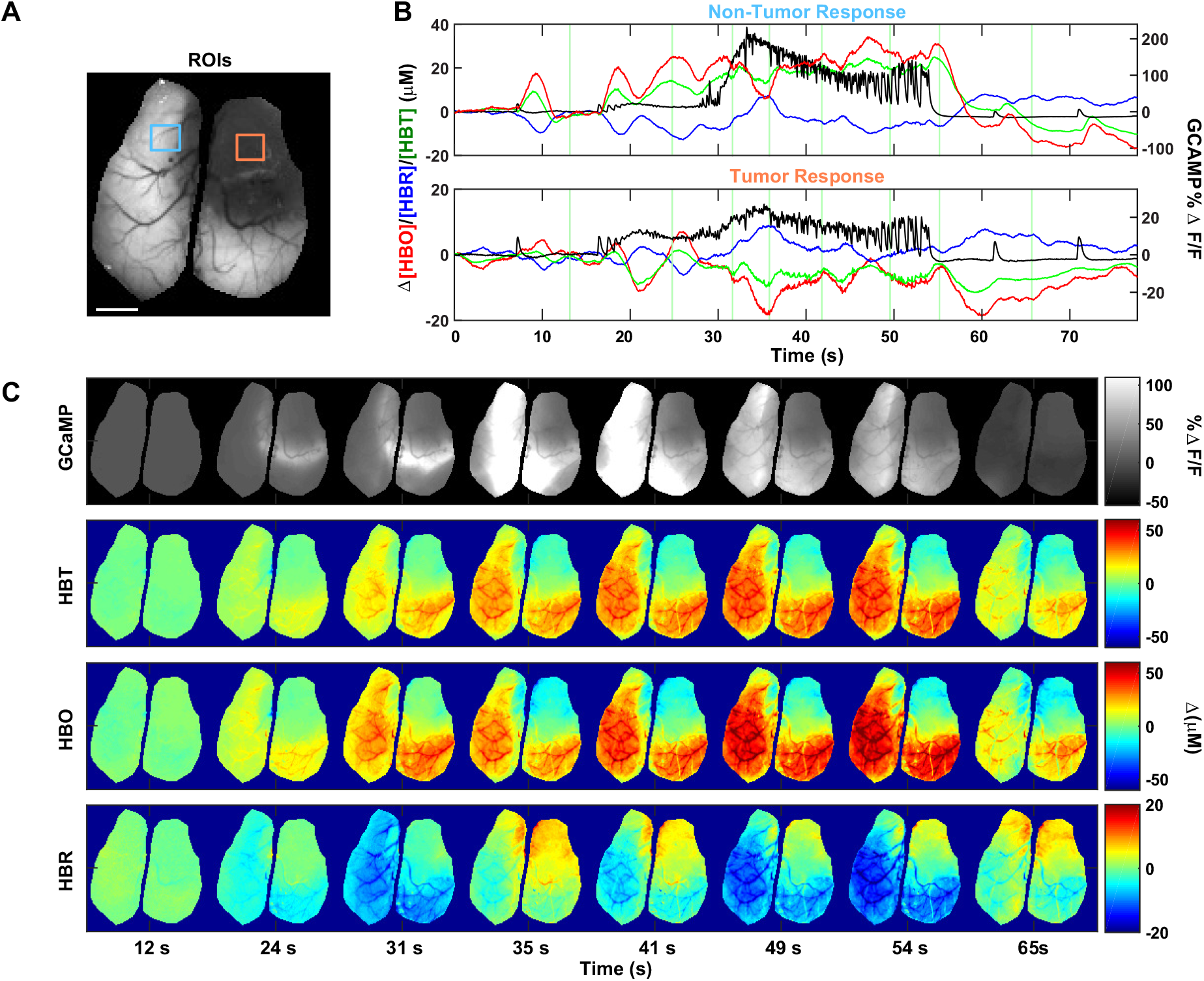
Neuronal and hemodynamic activity during a generalized seizure (Mouse 2, 34 days post injection). (A) Raw fluorescence image marked with relevant regions of interest in non-tumor (blue square) and tumor (orange square) regions. (B) Time series of a generalized seizure, shown for both GCaMP6f and hemodynamics and averaged across non-tumor and tumor regions. Timings of relevant frames of interest marked with vertical green lines. (C) Spatial patterns of GCaMP6f and hemodynamic data shown for relevant frames of interest during seizure event. Full seizure shown in Supplementary Video S4.

**Figure 7.**
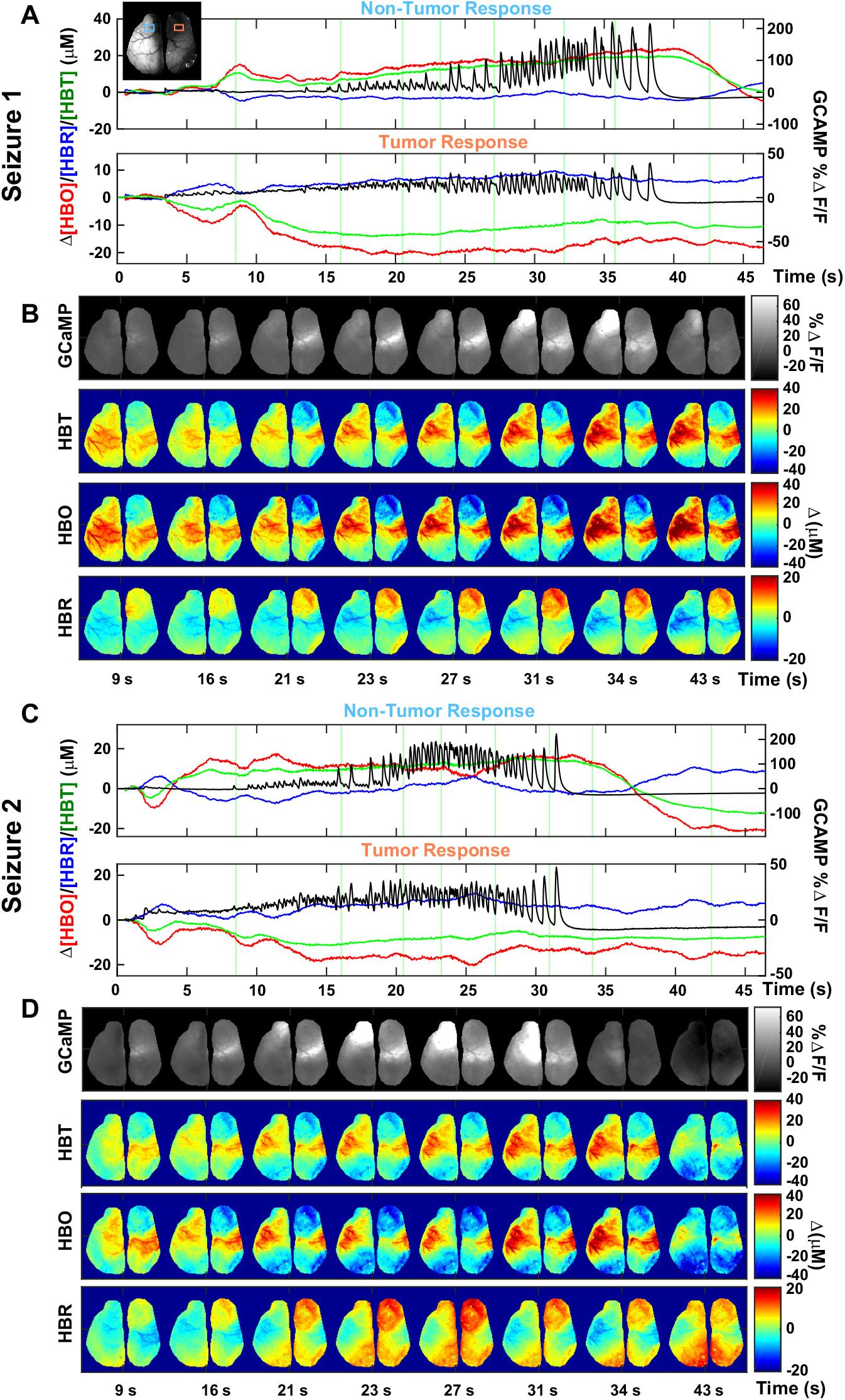
Neuronal and hemodynamic activity during two separate generalized seizures in mouse 1 (35 days post injection). (A,C) Time series of nonconvulsive generalized seizure, shown for both GCaMP6f and hemodynamics and averaged across non-tumor and tumor regions. Timings of relevant frames of interest marked with vertical green lines. Raw fluorescence image marked with relevant regions of interest in non-tumor (blue square) and tumor (orange square) regions on upper left. (B,D) Spatial patterns of GCaMP6f and hemodynamic data shown for relevant frames of interest during seizure event. Top seizure is shown in Supplementary Video S5.

Neuronal and hemodynamic signals extracted from the tumor and contralateral cortex are plotted in Figures 6 and 7, while Supplementary Figure S7A-B shows signals as a function of distance from the tumor center from anterior to posterior cortex for the seizure shown in Figure 6 and Supplementary Video S4. Examining hemodynamics associated with this seizure activity, in non-tumor-bearing cortex clear positive coupling to initial IIEs is visible, with large increases in HbT and HbO and decreases in HbR, as observed for both resting state neuronal activity and non seizure-initiating IIEs above. However, as neuronal events become more rapid and then further intensify, we note that the hemodynamic response does not exhibit a proportional increase. To test the linearity of the vascular response, a canonical hemodynamic response function was derived by averaging the HbT response from the four IIEs that occurred at the start of the seizure. This response was then convolved with the GCaMP6f fluorescence signal for the whole seizure. As shown in Supplementary Figure S7C, a linear hemodynamic response would predict an almost 10 times higher peak HbT response than observed. We infer that the maximum amplitude of the hemodynamic response to seizure activity, even in non-tumor bearing cortex, is limited by the maximal possible dilation of cortical vessels. A gradual decrease in HbO and increase in HbR during this ‘HbT saturation’ is consistent with oxygen consumption outstripping oxygen delivery suggesting widespread tissue hypoxia at the height of the seizure.

Examining hemodynamics within the tumor-burdened region for the same seizure event (Figure 6), we see an altered, attenuated and delayed hemodynamic response to the initial IIEs, consistent with IIE results shown in Figure 5, but with an even more pronounced post-event undershoot in HbT. Using the same method as above, convolution of this altered hemodynamic response function with neuronal activity in the tumor region during the seizure actually predicts a negative HbT response (hypoperfusion) to the seizure activity that quite closely matches measured HbT, without response-saturation effects (Supplementary Figure S7C-D). However, this ‘inverted coupling’ combined with the metabolic demand of the seizure activity appears to drive a clear decrease in HbO and an increase in HbR, suggesting significant hypoxia within the tumor region during seizure activity. These trends in both the tumor and non-tumor regions were recapitulated in both of the other seizure events recorded (shown in Figure 7 and Supplementary Video S5). These results suggest that the combination of glioma-induced seizures and region-specific disrupted neurovascular coupling results in significant hypoperfusion and hypoxia of the tumor tissue throughout the duration of tumor-induced seizure activity.

## Discussion

This study used wide-field optical mapping of both GCaMP6f and hemodynamics in awake mice at multiple time points during the progression of a diffusely infiltrating glioma. Our analysis revealed a wide range of changes in both neuronal activity and hemodynamics in the cortex that were associated with tumor progression, including disruptions in the synchrony of both neuronal and hemodynamic activity across the brain, alterations in the coupling between neuronal activity and blood flow, late onset of tumor-associated IIEs and eventually spontaneous seizures. These results demonstrate the power of WFOM, in combination with modern transgenic fluorescent labeling techniques, to provide new insights into the functional progression of brain diseases in-vivo. As a platform, WFOM could be used in other models of neurological disease to assess the interplay between cortical activity and disease progression. The method could also be used to evaluate the effects of new therapies for tumor suppression as well as anti-seizure measures on neuronal activity and vascular coupling. The multi-modal neural and hemodynamic read-outs of WFOM provide a further opportunity to interpret and predict hemodynamic signatures that might be detectable in humans using fMRI.

### Glioma progression leads to decorrelation of neuronal activity, IIEs and seizures

The changes in excitatory neural activity visualized during glioma progression began with attenuation of detectable neural activity which corresponded to a decrease in the density of neurons expressing GCaMP6f within the tumor. There was also a progressive desynchronization of neuronal activity in contralateral brain regions. The cause of desynchronization in the tumor compared to contralateral cortex is unclear, and could correspond to the onset of additional, spurious activity within the tumor, changes in the local generation of events due, perhaps, to metabolic or inhibitory / excitatory balance, but could also represent a disruption of connectivity between cortical regions (Bosma et al., 2008).

Tumor progression was accompanied by the onset of IIEs, which were captured by WFOM as events with characteristic temporal profiles of GCaMP fluorescence and hemodynamics, but which varied substantially in terms of apparent site of initiation, spatial spread and repetition frequency. Despite the lack of overt behavioral manifestations associated with IIEs, their increasing frequency during glioma progression suggests that IIEs may be more common in tumor patients than previously appreciated. Interictal epileptiform events have long been known to be highly specific, subclinical markers of epilepsy. While the frequency and temporal patterns of interictal discharges do not correlate well with seizure emergence in focal epilepsy syndromes in general, our data here suggest that increasing frequency of interictal discharges may correlate with, and perhaps even contribute to, the progressive neurological decline seen in glioma patients. Further studies of the potential clinical utility of interictal discharges as a biomarker of tumor progression may be warranted, as well as laboratory investigations into potential interventions aimed at reducing the IIE burden.

Three complete, spontaneous generalized seizures were also captured during in-vivo imaging, occurring at 34 and 35 DPI in two different mice. While the small size of our cohort limited our ability to access seizure frequency, the fact that these spontaneous events were seen during relatively short imaging sessions suggests that many more could be occurring during the late stages of disease. Notably, all of the generalized seizures that were seen during WFOM were immediately preceded by bursts of high amplitude discharges that appeared to begin at the infiltrative margin of the tumor, and then spread to surrounding cortex. This finding is consistent with electrophysiological data from ex vivo slice culture studies as well as clinical observations, suggesting that glioma-induced seizures arise from the infiltrated peritumoral cortex (Senner et al., 2004) (Pallud et al., 2014).

Together, these observations reveal a heterogeneous landscape of functional changes and interactions that occur during glioma progression, including widespread alterations in cortical neural activity leading to interictal discharges, and more severe focal alterations in and around the tumor that may predispose the brain to seizure activity. Marked histopathological findings at the tumor and tumor-margins included infiltration of glioma cells, neuronal loss, reactive gliosis and vascular changes. These finding implicate the glioma margin as an important place for continued functional characterization to understand the molecular and cellular basis for its role in initiation of epileptogenic activity.

### Progressive disruption of neurovascular coupling

Analysis of neural and hemodynamic WFOM data provided clear evidence for glioma-related disruption of neurovascular coupling within tumor-burdened regions of the cortex. This result is consistent with prior observations of significant interactions between infiltrating glioma cells and many aspects of the neurovascular unit and the microvasculature itself (Farin et al., 2006) (Watkins et al., 2014). Stimulus-evoked hemodynamics in the whisker region, which was not infiltrated by glioma, served as a control to show that normal-appearing hemodynamic responses could still be evoked throughout disease progression. However, analysis of coupling to spontaneous ‘resting state’ activity revealed profound systematic changes in coupling in regions affected by the tumor. Such changes in coupling of hemodynamics to neural activity could impact the reliability of pre-surgical fMRI measurements used to map functional regions proximal to the tumor boundaries (Zaca et al., 2014) (Sakatani et al., 2003), but could also potentially serve as biomarkers for disease progression if fMRI analysis can allow for regional alterations in the hemodynamic response function.

In regards to resting-state fMRI, our results suggest that both neuronal desynchronization and neurovascular disruption could contribute to apparent changes in fMRI-based measures of ‘functional connectivity’ related to tumor progression (Hadjiabadi et al., 2018) (Dierker et al., 2017). Nevertheless, these results do suggest a firm physiological basis for studies which have noted the ability of resting state fMRI data to delineate glioma tumor boundaries (Bowden et al., 2018) (Chow et al., 2016) (Agarwal et al., 2016).

Analysis of neurovascular coupling to IIEs in cortical regions outside the tumor revealed strong positive coupling (functional hyperemia), while the hemodynamic response was attenuated, delayed and distorted with an initial decrease in HbT in tumor-burdened regions. This response signature provided independent verification of altered coupling determined from resting state deconvolution analysis, and underscores that many aspects of dynamic physiology within the tumor region are altered. Our ability to examine these interactions in-vivo from the onset of IIEs onwards suggests that this imaging platform could be used to identify methods for reducing IIEs, or understanding their innate effects on behavior, cognition and neural dysfunction in the context of glioma progression.

Our observations of neurovascular dynamics during spontaneous generalized seizures also revealed marked differences in neurovascular coupling between tumor-bearing and non-tumor-bearing cortex. Even in non-tumor-bearing cortex with relatively normal coupling, our observations suggested rapid saturation of the hemodynamic response, while the amplitudes of neuronal activity greatly exceeded normal levels (>100% change in fluorescence) for sustained periods of over 30 seconds. This condition is likely to result in transient seizure-evoked hypoxia, consistent with studies exploring pharmacologically-induced seizures (Ma et al., 2013; Zhao et al., 2009). These brain-wide effects of glioma-induced seizure activity could be a major contributor to the increased morbidity and cognitive decline seen in low-grade glioma patients with intractable seizures (Klein et al., 2003; Ruda et al., 2010).

Within the tumor-bearing cortex, altered coupling had the exacerbating effect of decreasing perfusion of the region during seizure activity. This prolonged hypoperfusion, combined with the marked increase in neuronal activity and associated increase in metabolic demands, should result in severe tissue hypoxia within the tumor-bearing cortex. It is well established that tissue hypoxia can contribute to glioma progression through multiple mechanisms, including stimulating pathological vascular proliferation, production of tumor-promoting growth factors, and invasion into surrounding brain tissue (Cavazos and Brenner, 2016) (Hardee and Zagzag, 2012; Monteiro et al., 2017). In this context, our findings suggest that episodes of prolonged seizure-induced hypoperfusion of glioma infiltrated tissue could actually hasten progression of the disease.

These findings add to the growing body of evidence suggesting a bidirectional relationship between alterations in cortical activity and glioma growth and progression (Buckingham et al., 2011) (Campbell et al., 2012) (John Lin et al., 2017) (Johung and Monje, 2017) (Gillespie and Monje, 2018; Venkatesh et al., 2017) (Tewari et al., 2018). This work demonstrates the potential of WFOM in-vivo optical imaging in mice to improve our understanding of these pathophysiological phenomena, while providing a valuable new platform to discover and evaluate new preventive and protective treatments for the neurological consequences of glioma.

## Acknowledgments.

Two-photon images were collected in the Confocal and Specialized Microscopy Shared Resource of the Herbert Irving Comprehensive Cancer Center at Columbia University, supported by NIH grant #P30 CA013696. The two-photon microscope was purchased with NIH grant #S10 RR025686. Assistance with breeding, animal care, data acquisition and analysis was provided by members of the Hillman Lab including Carla Kim, Roni Moon and Ericka Wu. This study built upon foundational work by Mariel Kozberg, Sasha Rayshubskiy, Sarah DeLeo and Brenda Chen and collaborators Charles Mikell, Angel Lignelli, Guy McKhann and Mark Otten. This work was supported by grants R01NS076628 (Hillman), R01NS063226 (Hillman), RF1 MH114276 (BRAIN, Hillman), CTSA pilot funding from NCATS UL1 TR000040 (Hillman / Canoll), Columbia ROADS grant RG31 (Hillman / Zheng), R03 NS090151-01 (Canoll), James F. McDonnell Foundation BTEC Award (Canoll /Gutmann, /Ellisman,), and American Epilepsy Society Pilot Study Grant (Canoll/Schevon).

## Author Contributions

Conception and design; EMCH, PC and SHK. Development of methodology; EMCH, PC, SHK, AD, THZ, DNT, MS, YM. Investigation and data acquisition; SHK, AD, KP, THZ. Data analysis and interpretation; MKM, SHK, EMCH, AD, MS, JG, CS. Technical and material support; AM, NH, JG, CS, DSC, EMCH. Writing and revision of the manuscript; MKM, SHK, AD, EMCH and PC.

## Materials & Methods

Please see Table 1 for a complete list of materials and software used in this study.

**Table 1.**

| REAGENT or RESOURCE | SOURCE | IDENTIFIER |
| --- | --- | --- |
| **Antibodies** |  |  |
| rabbit anti-Aquaporin 4 | Millipore Sigma | HPA014784 |
| rat anti-HA tag | Millipore Sigma | 11867423001 |
| rabbit anti-NeuN | Cell Signaling Technology | 12943 |
| rabbit anti-CD34 | Abcam | ab81289 |
| Goat anti-rat Alexa Fluor 568 Conjugate | Life Technologies | A11077 |
| Goat anti-rabbit Alexa Fluor 633 Conjugate | Life Technologies | A21071 |
| Goat anti-rabbit Alexa Fluor 568 Conjugate | Life Technologies | A11036 |
| **Chemicals, Peptides, and Recombinant Proteins** |  |  |
| Isolectin GS-IB<sub>4</sub>, biotin-XX Conjugate | Life Technologies | I21414 |
| Streptavidin, Alexa Fluor 633 Conjugate | Life Technologies | S21375 |
| Quadrol | Millipore-Sigma | 122262 |
| Urea | Millipore-Sigma | U4883 |
| N-butyl-diethanolamine | Millipore-Sigma | L09953 |
| Antipyrine | Fisher Scientific | 104971000 |
| Nicotinamide | Fisher Scientific | 128271000 |
| Triton X-100 | Fisher Scientific | BP151 |
| **Experimental Models: Cell Lines** |  |  |
| Mouse p53<sup>-/-</sup> PDGFA<sup>+</sup> mCherry-Luciferase<sup>+</sup> RPL22<sup>HA</sup> | This study | N/A |
| **Experimental Models: Organisms/Strains** |  |  |
| C57BL/6J-Tg(Thy1-GCaMP6f)GP5.17Dkim/J | Jackson Laboratory | 025393 |
| **Software and Algorithms** |  |  |
| FIJI/ImageJ | NIH | https://imagej.net/Fiji |
| NIS Elements AR v5 | Nikon | https://www.microscope.healthcare.nikon.com/products/software/nis-elements/nis-elements-advanced-research |
| Zen (Blue edition) | Zeiss | https://www.zeiss.com/microscopy/us/products/microscope-software/zen.html |
| Prism, Version 7 | GraphPad | https://www.graphpad.com/scientific-software/prism/ |
| MATLAB 2016a – 2018b | MathWorks | https://www.mathworks.com/products/matlab.html |
| Modified Beer-Lambert Conversion<br>Excitation-Emission Hemodynamic<br>Correction | (Ma et al., 2016a) | https://www.ncbi.nlm.nih.gov/pmc/articles/PMC5003860/ |
| Deconvolution Resting State Analysis | (Ma et al., 2016b) | https://www.pnas.org/content/113/52/E8463 |
| Interictal event identification via<br>prominence and full-width at half<br>prominence | (Steinmetz et al., 2017) | https://www.ncbi.nlm.nih.gov/pmc/articles/PMC5604087/ |

### Glioma model and preparation of Thy1-GCaMP6f mice for imaging

All procedures were reviewed and approved by the Columbia University Institutional Animal Care and Use Committee (IACUC).

The primary diffusely infiltrating gliomas were generated by injecting PDGFA-IRES-Cre expressing retrovirus into the subcortical white matter of transgenic C57BL/6 mice that harbor floxed p53, stop-flox RPL22^HA^, and stop-flox mCherry-luciferase. The resulting retrovirus-induced tumors show the histological features of diffusely infiltrating gliomas (Mela, Torres, Canoll, unpublished). The mouse glioma cells were isolated from a retrovirus-induced tumor and expanded in culture as previously described (Sonabend et al., 2013). Diffusely infiltrating gliomas were induced in Thy1-GCaMP6f mice via orthotopic transplantation of these mouse glioma cells as detailed below.

Thy1-GCaMP6f mouse littermates (Dana et al., 2014) were weaned at postnatal day 21 and housed together in temperature and humidity controlled facilities with 12-hour light/dark cycles and ad libitum water and food. Adult (∼2.5 months old) Thy1-GCaMP6f positive mice (C57BL/6J-Tg(Thy1-GCaMP6f)GP5.17Dkim/J, The Jackson Laboratory Stock No. 025393) were acclimated to human handling and running wheel. Following acclimation, mice were anesthetized using isoflurane (3-4% for induction, 2% for maintenance), placed in a stereotaxic frame with ear bars (Kopf) on top of a homeothermic heat pad (temperature maintained at 37°C) and injected with burprenorphine analgesic (0.05 mg/kg). An incision was made and scalp overlying the superficial cortical area excised. The dorsal skull surface was thinned to transparency using a dental drill (Aseptico) with a .9 mm stainless steel burr (Fine Science Tools) while irrigating often with sterile 0.9% saline. Immediately after skull thinning a small hole was created using a 20g needle and glioma cells were injected into the subcortical white matter of right frontal cortex (stereotaxic coordinates relative to bregma= 2.0mm anterior, 2.0mm lateral, 1.5 mm deep) using a Hamilton syringe with a 33g needle (5×10^4^ cells in 1 μl at a flow rate of 0.25 μl/min).

The skull was allowed to dry and liquid cyanoacrylate glue was applied in a thin layer, simultaneously covering the injection hole. The bordering skin was sealed with cyanoacrylate tissue adhesive and gel cyanoacrylate glue was used to bond a custom laser-cut acrylic plastic head plate to the border of the skull. Silicone elastomer was applied to the surface to further protect the thinned-skull preparation during recovery and reapplied between imaging sessions. Implanted animals were housed in separate cages and allowed to recover for 72 hours with continued buprenorphine analgesic administration and twice-daily monitoring.

After recovery, in-vivo imaging (details below) was conducted 3-5 times a week for up to 36 days post-injection. Animals were head restrained for 15-30 minute sessions in identical restraint hardware separate from the imaging system for the first week of imaging for better acclimation. Animals were monitored daily for symptoms of tumor burden. If animals exhibited symptoms of pain, altered behavior or weight loss of 20% or more, animals were euthanized via cardiac perfusion under 5% isoflurane with PBS and brains were extracted and fixed in 10% neutral buffered formalin.

### Functional imaging Data acquisition

WFOM was conducted to collect simultaneous reflectance and GCaMP6f fluorescence signal as previously described in (Ma et al., 2016a), using high-speed strobed LED illumination (490, 530 and 630 nm) time-locked with an sCMOS camera (Andor Zyla) at 50.25 Hz. Reflectance and fluorescence emission light from the brain was collected with a 500-640 nm bandpass filter to reject illumination wavelengths and focused using an AF Micro-NIKKOR 60 mm lens (Nikon). Webcam monitoring of the mouse’s behavior was collected throughout imaging with infrared illumination.

Mice were imaged longitudinally throughout the period of tumor development, starting at 4 days post glioma cell injection, with each mouse having at least 8 imaging sessions. Each animal was head-restrained for a maximum of two hours per imaging session. Resting state recordings involved 12 to 18 recordings (180 s duration each) over the course of each imaging session. For whisker barrel stimulation, recordings were 30 s in duration (10 s pre-stimulation, 5 s whisker stimulation, 15 s post-stimulation). A total of 30 recordings were performed for each of the right and left whiskers per imaging session.

### Analysis

#### Conversion of raw data

Reflectance signal from red (630 nm) and green (530 nm) illumination were converted using the modified Beer-Lambert method and known hemoglobin absorption spectra into changes in oxygenated (HbO) and deoxygenated hemoglobin (HbR) concentrations. Total hemoglobin (HbT) was calculated as the sum of HbO and HbR. Fluorescence signal (excited at 490 nm) was converted to ΔF/F (mean normalized) and corrected for hemodynamic contamination using the Excitation-Emission (‘Ex-Em’) method as described in (Ma et al., 2016a) which estimates excitation and emission attenuation of the fluorescent light using the simultaneously measured dynamics of HbO and HbR. Excitation and emission pathlength coefficients were estimated according to optimal fluorescence signal correction, and coefficients permitted to change during the course of tumor progression due to possible changes in optical properties of the cortex.

After conversion, data for each mouse was registered to the field of view of its first recording session. Epochs where the animal was neither walking nor running were identified using a thresholded standard deviation of adjacent webcam frames and used for subsequent functional connectivity and deconvolution analysis. Data was corrected for minor flickering from illuminating LEDs by multiplication with the low-pass filtered full image mean time course and division by the mean full image time course.

#### Functional connectivity analysis

K-means clustering (Pearson Correlation distance, k=22) was applied to the first session GCaMP6f resting state dataset to identify functional regions of interest. Pearson correlation analysis was then applied pixel-by-pixel to both GCaMP6f and HbT data following preconditioning with a 5×5 pixel spatial box filter. Functional regions (glioma injection site, contralateral to glioma, forepaw, whisker barrel and visual cortex) were selected from k-means clusters.

#### Whisker stimulus averaging

Whisker stimulus time courses extracted from the localized region of response were averaged per session. Trials containing running during, immediately before, or after the stimulus period were identified by thresholding the standard deviation from images from corresponding webcam data and were excluded from averages. Trials where hemoglobin concentrations continued to drift post-stimulus rather than returning to baseline were assumed to have been immediately followed by running and were also excluded from averages.

#### Hemodynamic response function deconvolution

Data was examined and sorted to isolate epochs of recordings that were at least 30 seconds in duration and did not include either running or IIE or seizure activity. Raw fluorescence, red and green reflectance signals were then extracted from 8 regions of interest across the cortex, with data from each day and session registered to each other to ensure consistent region selection. Owing to the sensitivity of deconvolution to noise and filtering, and higher levels of noise on red signals compared to green, deconvolution was performed using simplified pre-processing consisting of 1) conversion of all signals to % change using the average of 6 seconds of initial baseline, 2) correction of fluorescence for hemodynamics by dividing by the green reflectance signal raised to the power 1.3 corresponding to an estimate of the pathlength difference between fluorescence and diffuse reflectance and validated by careful inspection to ensure that corrected fluorescence did not follow a clear trend that matched green reflectance, and 3) high-pass temporal filtering of all signals at >0.05 Hz to remove slow drifts, and low pass filtering of green and red signals to reduce noise at <1 Hz. Deconvolution was then performed using a regularized convolution matrix approach in Matlab™ (Ma et al., 2016b). A 3.6 delay was introduced between the GCaMP and red or green traces prior to deconvolution to allow the output to include prediction of pre-neural event dynamics. For every trial, the correlation between the original raw red or green data, and the result of convolution of the HRF solution with the original corrected fluorescence trace was calculated and is depicted in plots shown in Figure 3C and Supplementary Figure S4C and G, with the correlation coefficient for each trial shown as a dot, each horizontal bar depicting the mean for that imaging session, and error bars showing the standard deviation. Deconvolution resulted in HRFs for each trial for both green and red reflectance, which were then converted to equivalent changes in HbO, HbR and HbT. Plots in Figure 3B and Supplementary Figure S4B and F show individual HRFs for every trial as thin lines, with a bold line representing the mean average response time-course for that region and measurement day. Colormaps in Figure 3D and Supplementary Figure S4D and H depict the amplitude of the HRF on the colorscale indicated, and depict the mean HRF for each imaging session and region of interest. This analysis was repeated with a wide range of different deconvolution parameters, denoising methods and correction approaches to reduce the effects of low signal to noise in later GCaMP6f signals (caused by neuronal loss) and spurious noise in red reflectance as well as potential changes in the validity of the hemodynamic correction within the growing tumor. Results shown are representative of the majority of outputs from this repeated analysis.

#### Interictal event analysis

Onset analysis shown in Figure 4D began with finding the peak time of each IIE from a representative GCaMP time-course. Data in this case was acquired at 25.8 Hz. For each event, a 0.3 second epoch (8 time-points) spanning the peak was examined for every pixel, after down-sampling the image to 64×64 pixels to improve signal to noise. A threshold corresponding to 20% of the maximum GCaMP value across the cortex at the peak time was chosen for each event. For pixels whose maximum within the event epoch exceeded this threshold, we calculated the time taken for the GCaMP6f signal to rise to this threshold value, improving precision by spline interpolating time-courses 10x to 258 Hz. The resulting colormap for each event depicts time taken for each locations to reach this GCaMP threshold, with the minimum onset time value within the map subtracted such that the earliest location at t = 0 shows the site of first onset. Plots were generated using the contourf function in Matlab™. This onset time analysis was also repeated using 50% of the peak value of each pixel’s signal for each event as the threshold (which normalizes for the amplitude of the event at each location). Onset maps in this case had the same general pattern and sites of localization.

To generate the plot shown in Figure 4E, the total number of IIEs identified per session was normalized by the total duration of resting state recordings acquired in minutes to represent changes in IIE frequency observation over the course of tumor development.

For the analysis shown in Figure 5, spike triggered averaging analysis of hemodynamic signals was applied to interictal events (IIEs) observed in resting state GCaMP6f data. The findpeaks Matlab™ function was first applied to GCaMP6f data, with events then user-verified to be IEE-type spikes of at least 2% (but typically closer to 10% or more) ΔF/F amplitude and at least .1 s in width/duration. Corresponding HbO, HbR and HbT signals were then time-aligned based on these IEE events and averaged. Peak amplitudes for GCaMP and hemodynamic signals averaged across all IIEs from all sessions for each IIE were calculated by finding the maximum (or in the case of [HBR], the minimum) value and time to half peak, for each mouse as shown in Panel C of Figures 5 & S6 for each event. Time to half peak was determined by finding the location of the first occurrence of half the previously found peak amplitude in the signal.

Further analysis of IIEs again utilized the findpeaks MATLAB function where potential events. Usage of prominence and full-width at half-prominence of peaks was incorporated from methods from (Steinmetz et al., 2017). For optimal separation from noise and physiological occurrences of GCaMP6f activity, IIE’s were constrained to have a minimum peak of 7%, minimum prominence of 2%, full-width at half-prominence between .1 s and 1 s, a prominence to full-width ratio greater than twice the ratio of the total data peak prominences median to total data peak widths median, and at least 10 frames away from the nearest interictal event. Each event was user-verified with time courses and data visualization to eliminate any spurious motion artifact related events. Event image series were thresholded at 6% fluorescence to detect events and observe spatial propagation. IIEs that propagated to involve more than 50% of the masked brain pixels were categorized as global interictal events, whereas those that did not propagate, or involved less than 50% of the cortex were categorized as localized interictal events (Figure 4 and Supplementary Figure S5).

#### Seizure analysis

Generalized seizure epochs were user-identified and signal time courses were extracted from selected regions of interest within and contralateral to the glioma injection site, as well as from the tumor margin. Predicted HBT signal for a given generalized seizure was estimated using the average hemodynamic response of preceding interictal events in the same region of interest as the original signal convolved with the region GCaMP6f time course.

### Histological analysis of brain sections

#### Preparation of brain sections for confocal microscopy

Histological analysis of free-floating sections was performed on 2 identically-prepared tumor bearing mice that did not undergo in vivo imaging (Figure 1 and Supplementary Figure S1A). Coronal sections of 40 μm thickness were obtained on a vibratome. Sections were washed in PBS prior to being delipidated and decolorized in sequential incubations in ½ -dH_2_O diluted CUBIC reagent 1A (10% w/w triton X-100, 5% w/w Quadrol, 10% w/w Urea, 25 mM NaCl; 30 minutes at room temperature), followed by incubation in complete Reagent 1A for 3.5 hours at 37°C with gentle shaking (Susaki and Ueda, 2016) (Pavlova et al., 2018). Following PBS washes, the sections were blocked in PBS containing 10% v/v Normal goat serum and 0.3% w/w Triton X-100. Primary antibody incubations were performed at 4°C for 72 hours followed by secondary antibody incubation for 6 hours at room temperature. Confocal images of CUBIC Reagent 1A clarified coronal brain sections were captured using 488 nm, 561 nm and 639 nm excitation on a Zeiss LSM 800 confocal microscope. Low-power scans of coronal sections were acquired using a 20x/0.75 NA objective. High-power images were acquired with a 40x/1.3 NA oil immersion objective. Maximum projections of Z-stacks were generated using FIJI/ImageJ.

#### Post mortem histological analysis of whole brain tissue by two-photon microscopy

Whole fixed brains of mice that underwent in vivo imaging were clarified using the CUBIC-cancer method as described (Kubota et al., 2017). Briefly, excised brains were delipidated in ½-dH_2_0 – diluted CUBIC-L solution (10w%/10w% N-butyldiethanolamine/Triton X-100) for 6 hours followed by incubation in complete CUBIC-L for 5 days with gentle shaking at 37°C. Brains were subsequently washed with PBS and blocked in 10% normal goat serum, 0.5w% Triton X-100 for 6 hours, followed by staining with the indicated primary antibodies (Supplementary Figure S1C-D) for 5 days at room temperature in blocking buffer. After multiple PBS washes, secondary antibody incubations were performed for an additional 4 days at room temperature in blocking buffer. Following final PBS washes, brains were immersed in CUBIC-R solution (45w%/30w% antipyrine/nicotinamide) and stored and imaged in the same solution. High-resolution images of CUBIC-cancer clarified post-mortem brains were acquired using a Nikon A1RMP multiphoton confocal microscope and a 25×/1.10 NA coverslip-corrected water immersion IR lens objective. For two-photon acquisition, a Chameleon II laser was tuned to 820 nm and images were captured on a spectral detector. Images were taken at 5 μm steps, unmixed and stacks were generated with NIS Elements AR or FIJI/ImageJ software.

## Supplementary Figures

**Supplementary Figure S1. (Related to Figure 1).**
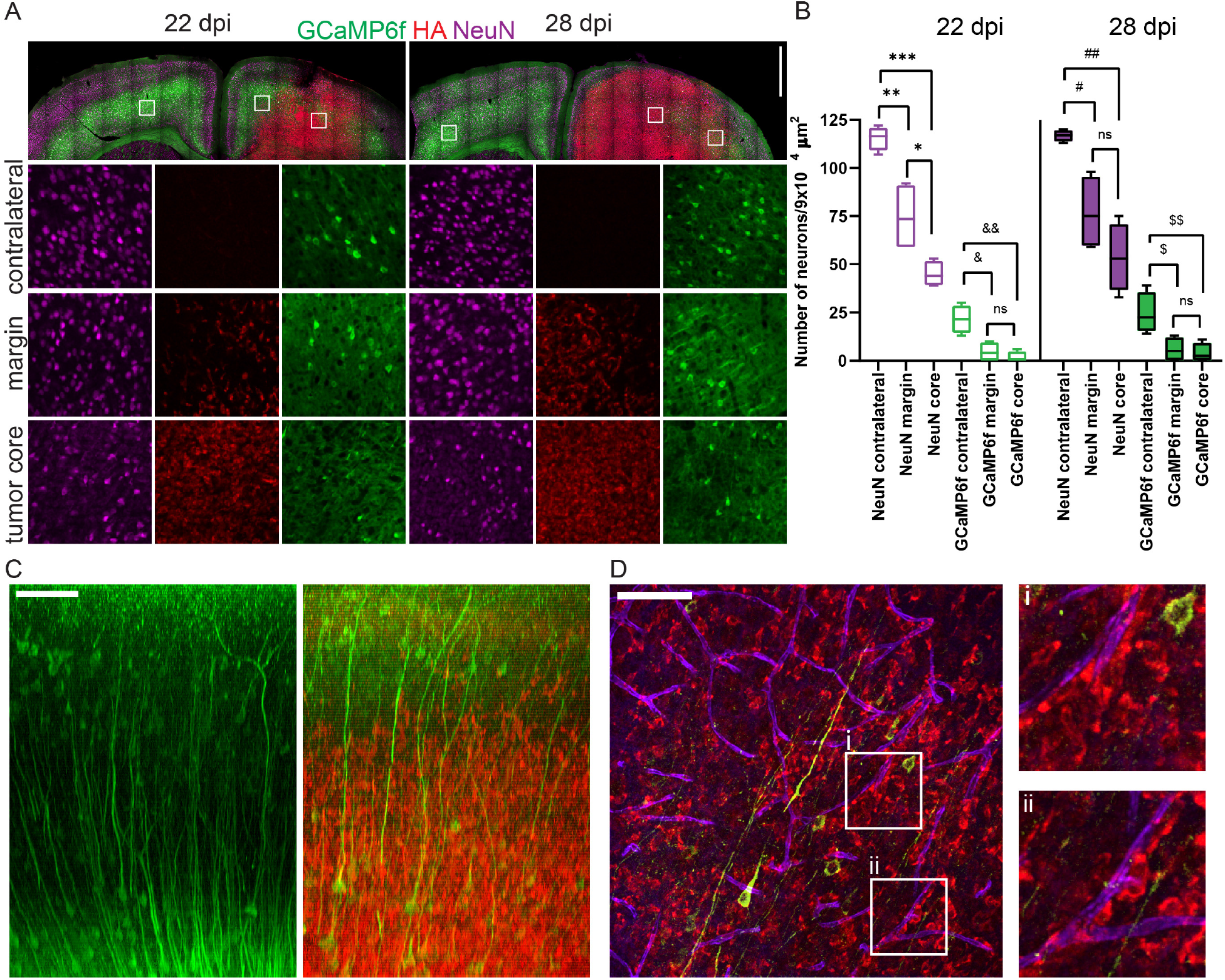
Loss of neurons, perineuronal satelitosis and perivascular invasion in the cortex of glioma-bearing mice. (A) Coronal sections obtained from GCaMP6f^+^ mice at 22 or 28 days post tumor cell injection stained with NeuN to identify neurons and HA for tumor cells. Stitched images show a single confocal plane. Bar, 1000 µm. Lower panels, enlarged insets. (B) Numbers of NeuN and GCaMP6f+ neurons from layer 5 (per 9×10^4^ μm^2^) in tumor margins, core and contralateral to the tumor. \**P*=0.02; \*\**P*=0.005; \*\*\**P*=0.00004; ^&^*P*=0.008; ^&&^*P*=0.002; ^#^*P*=0.006; ^##^*P*=0.0004; ^$^*P*=0.02; ^$$^*P*=0.01. Two-tailed, unpaired t-test was used to calculate significance. (C, D) Two-photon images of clarified post-mortem brain of mouse 1 reveal intermingling of tumor cells with neurons and peri-vascular patterns of invasion. (C) XZ projections of confocal volumes from contralateral (left) and ipsilateral (right) hemisphere of the brain from mouse 1, imaged post-mortem. Tumor cells (HA; red) invade the cortex and intermingle with GCaMP6f positive neurons (green). Bar, 100 μm. (D) Representative field from tumor margin showing GCaMP6f^+^ neurons (green), tumor cells (HA; red) and AQP4+ astrocytic end-feet (purple). Enlarged insets show perivascular association of glioma cells that closely associated with the abluminal surface of AQP4+ astrocyte end-feet. Bar, 100 μm.

**Supplementary Figure S2. (Related to Figure 2.).**
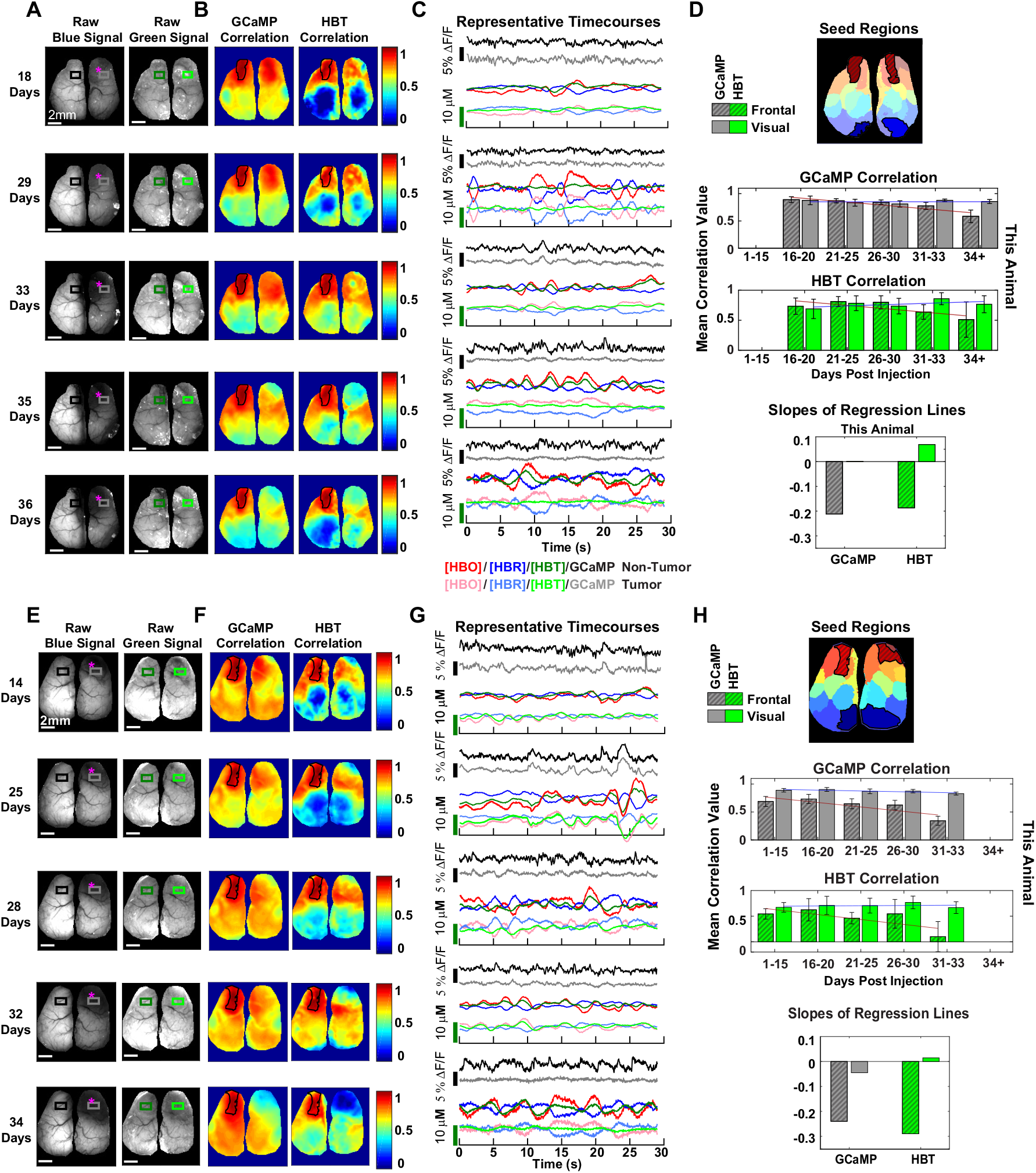
Correlation of GCaMP6f and hemodynamic signals between tumor and non-tumor regions during tumor progression (Mouse 1 (top) and 3 (bottom)) (A, E) Images of raw fluorescence signal showing progressive loss of GCaMP6f fluorescence with no corresponding loss of hemodynamic signal. Magenta asterisks indicate approximate site of glioma injection. (B, F) Maps of correlation to a seed region, outlined in black, for GCaMP6f and hemodynamic data.(C, G) Individual time courses taken from representative runs and averaged across regions in the tumor and non-tumor areas of the brain. (D, H) Top: Map of k-means seed regions used in correlation analysis with frontal (red diagonal hatch line fill). Center: Bar plots of both GCaMP6f and hemodynamic correlation in frontal (gray with diagonal hatch line fill) and visual regions (gray fill) as tumor progresses, shown for one animal. Bottom: Slopes of regression lines for GCaMP6f and hemodynamic correlation values shown for one representative animal and across all animals (error bars are standard error across animals). Significance across all animals was calculated using paired, two-sample, two-tailed t-tests between non-tumor and tumor regions for each channel at significance of p<0.05(*) and p<0.005 (**).

**Supplementary Figure S3. (Related to Figure 3).**
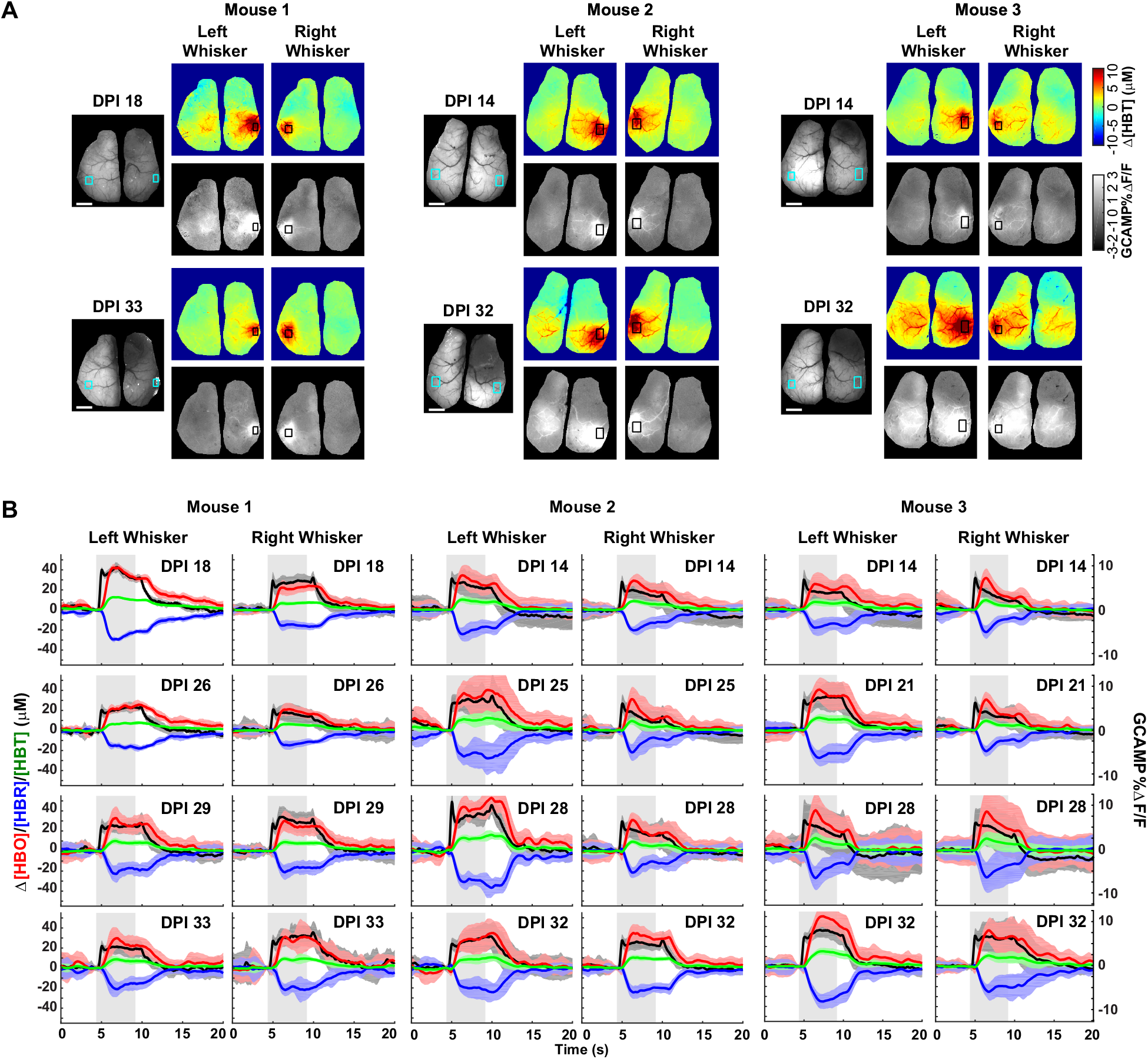
Whisker stimulus responses during tumor progression. (A) (Left) Raw fluorescence image showing location of whisker barrel in one early and one late day of imaging. (Right) Average total hemoglobin image and GCaMP6f image during the stimulus period for the same days as the fluorescence images. ROIs for the time courses below are marked with squares for all three images. (B) GCaMP6f and hemodynamic responses to 5 second left and right whisker stimulus for all animals over the course of tumor development (days post injection indicated on the upper right corner of each plot). Error bars indicate standard deviation across stimulus trials. Gray block indicates whisker stimulus time.

**Supplementary Figure S4. (Related to Figure 3).**
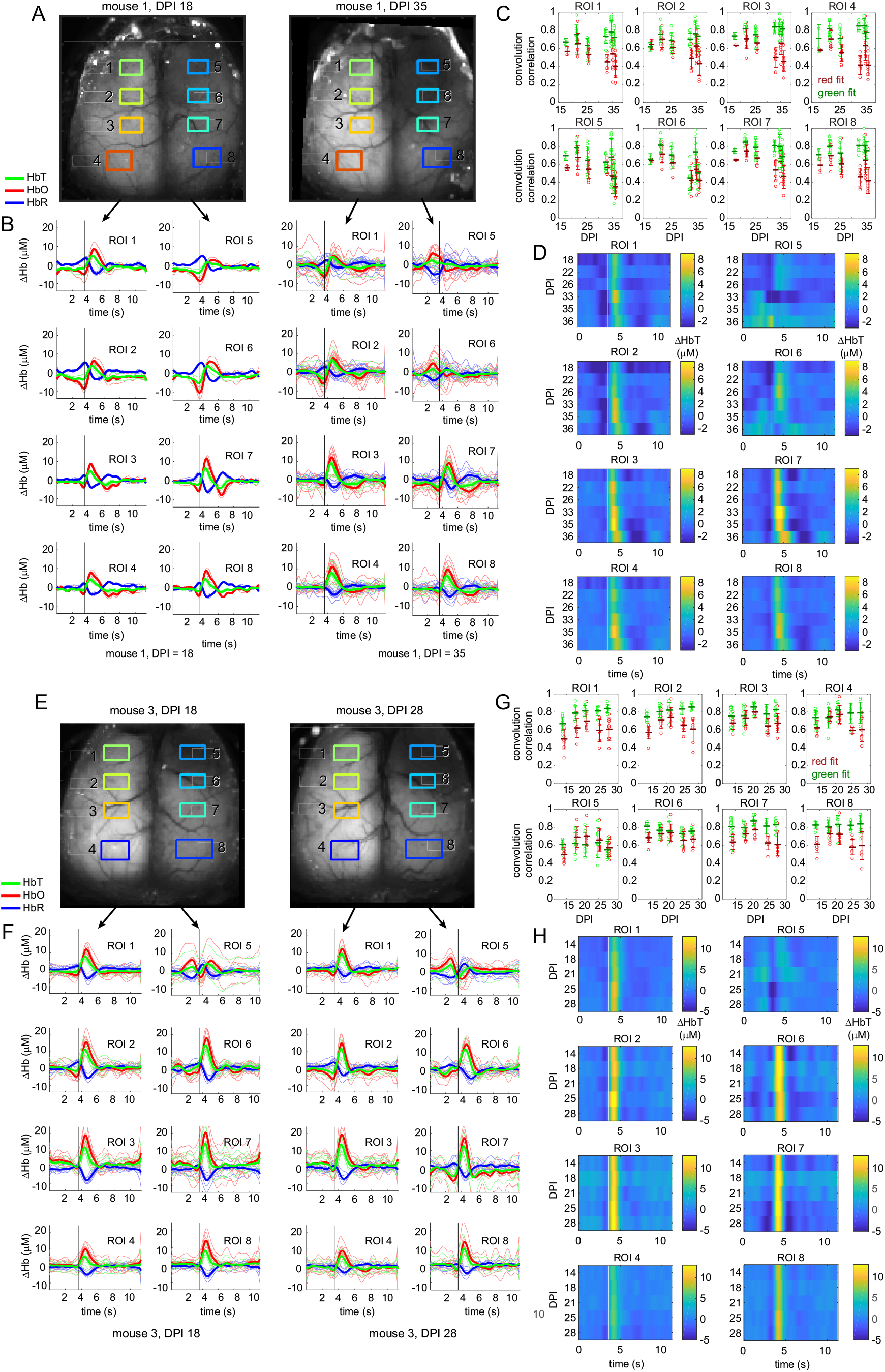
Analysis of regional changes in neurovascular coupling during tumor progression by deconvolution of resting state neural activity from hemodynamics in two additional mice. (A) and (E) show the brains of mice 1 and 3 respectively at early and late time-points with regions of interest spanning the cortex contralateral and ipsilateral to the tumor. (B) and (F) show hemodynamic response functions (HRFs) calculated using deconvolution of resting state epochs (see STAR methods), with fine lines showing individual trial results, and bold lines showing the average for each imaging session. The left two columns show ROIS 1-8 at 18 days past injection (DPI) showing bilaterally consistent responses in which HbT increases following the black line which denotes the peak of a spontaneous neural event. The second two columns show HRFs from the same 8 ROIs at 35 / 28 DPI. Here, the contralateral side has consistent hemodynamic responses, whereas the tumor-affected side shows profound changes including hyperemia prior to the event followed by an evoked decrease in HbT within the tumor, and an intermediate response at the tumor boundary where the neural event occurs during a distinct decrease in HbT. (C) and (G). Show correlation coefficients of each trial between the original red and green signals, and signals predicted by the convolution model. (see STAR methods). Circle = single run, bar = average over all trials, whiskers = standard deviation. Later days for tumor regions show degraded correlations suggesting worsening fits to a linear HRF fit. Lower correlation values for red compared to green reflectance correspond to worse noise on raw red signals. (D) and (H). Depicts aggregate average HRF results for all measurement sessions over the same 8 ROIs. The attenuation of the hemodynamic response within the gradually progressing tumor region is clear, along with the trends of initial low HbT and eventually pre-event hyperemia. The same analysis is shown for mouse 2 in Figure 3.

**Supplementary Figure S5. (Related to Figure 4).**
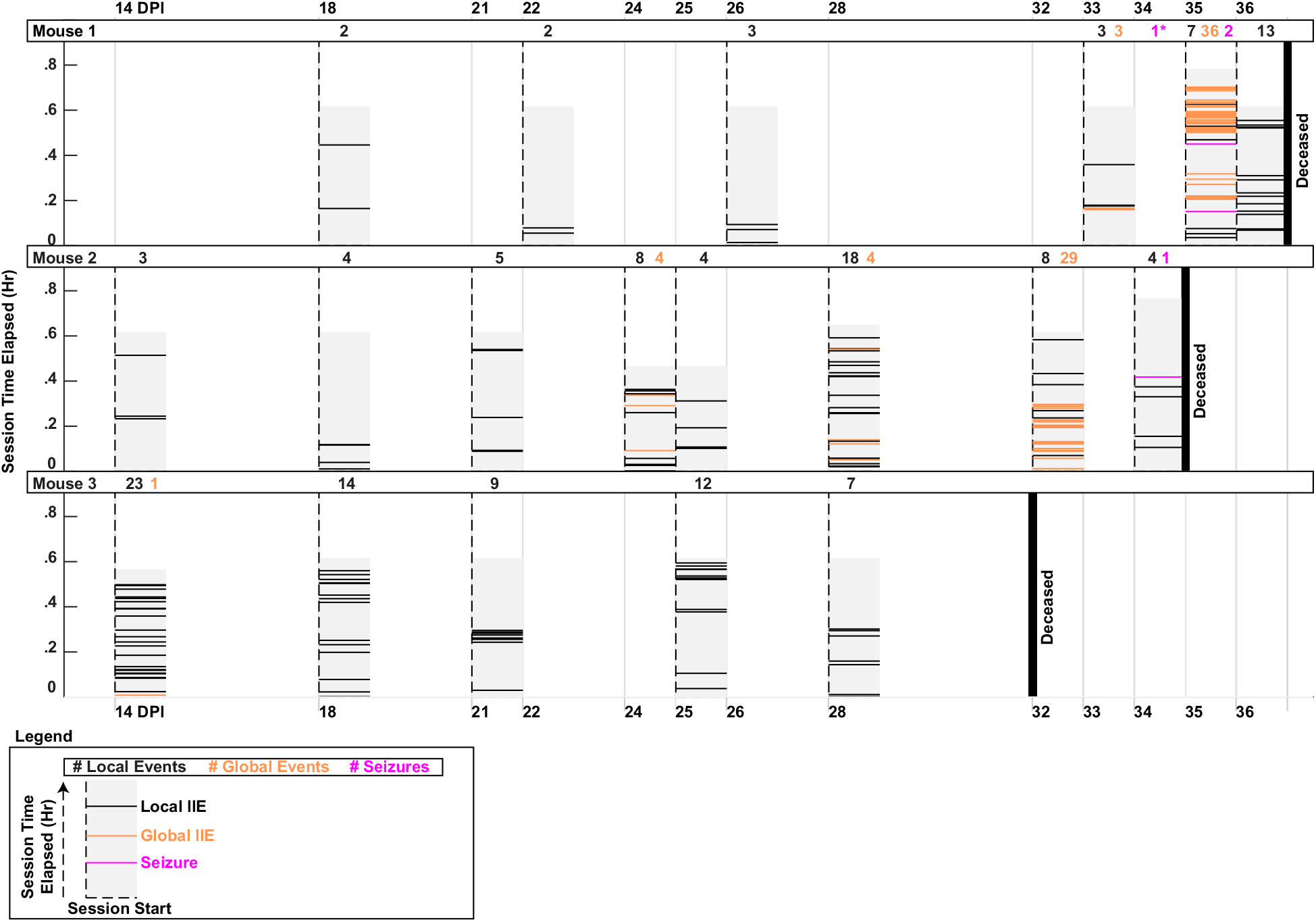
Interictal event frequency and timing over entire imaging period. Full timeline of occurrence of two types of interictal events, and seizures, in each animal during recording sessions (shaded in light gray blocks). The day of each recording session is indicated with a dotted vertical black line. Total duration of resting state parts of recording sessions varied from .4 hours up to .9 hours. Local events, global events and generalized seizures are indicated with horizontal black, orange and magenta lines. One convulsive generalized seizure, which was observed in Mouse 1 but not imaged, is indicated with magenta font and asterisk under 34 days post injection. Dark bold vertical lines indicate the day of death of animal.

**Supplementary Figure S6. (Related to Figure 5).**
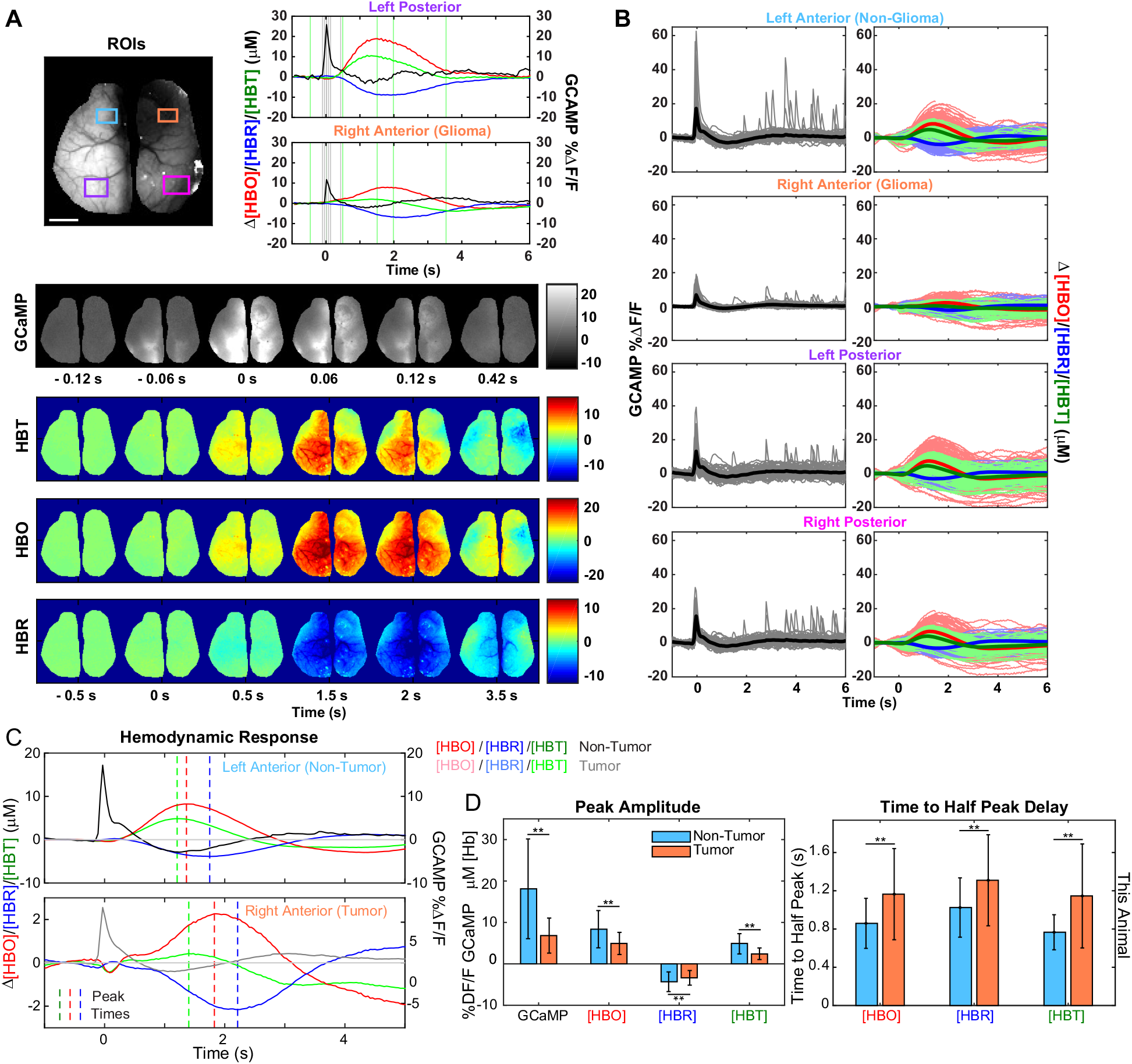
Region-specific alterations in neurovascular coupling during interictal events (Mouse 1) (A) Top Left: Raw fluorescence image marked with relevant regions of interest. Top Right: Time series taken from a representative event (33 days post injection) and averaged across pixels in the left posterior and right anterior (tumor) regions. Relevant frames of interest for GCaMP6f marked with gray vertical lines and for hemodynamics with green vertical lines. Bottom: Spatial patterns of GCaMP6f and hemodynamic data shown for relevant frames of interest during the event. (B) Temporally aligned time series for all events from all sessions (all days) in one representative animal averaged across pixels in various regions of interest and shown for both GCaMP6f and hemodynamic data, with the average plotted in bold. (C) Temporally aligned time series, averaged across all events, showing the hemodynamic responses in non-tumor and tumor regions, with vertical bars indicating peak times. The vertical axes for the GCaMP6f and hemodynamic right anterior (tumor region) time courses are scaled differently due to amplitude differences. (D) Bar plots of peak amplitude of GCaMP6f and hemodynamic data (left column) and time to half peak of hemodynamic data (right column), shown for one representative animal (top row) and across all animals (bottom row). Error bars are standard deviation across trials for one representative animal, and standard deviation across all animals. Significance across all animals was calculated using paired, two-sample, two-tailed t-tests between non-tumor and tumor regions for each channel at significance of p < 0.05 (*) and p < 0.005 (**).

**Supplementary Figure S4. (Related to Figure 7).**
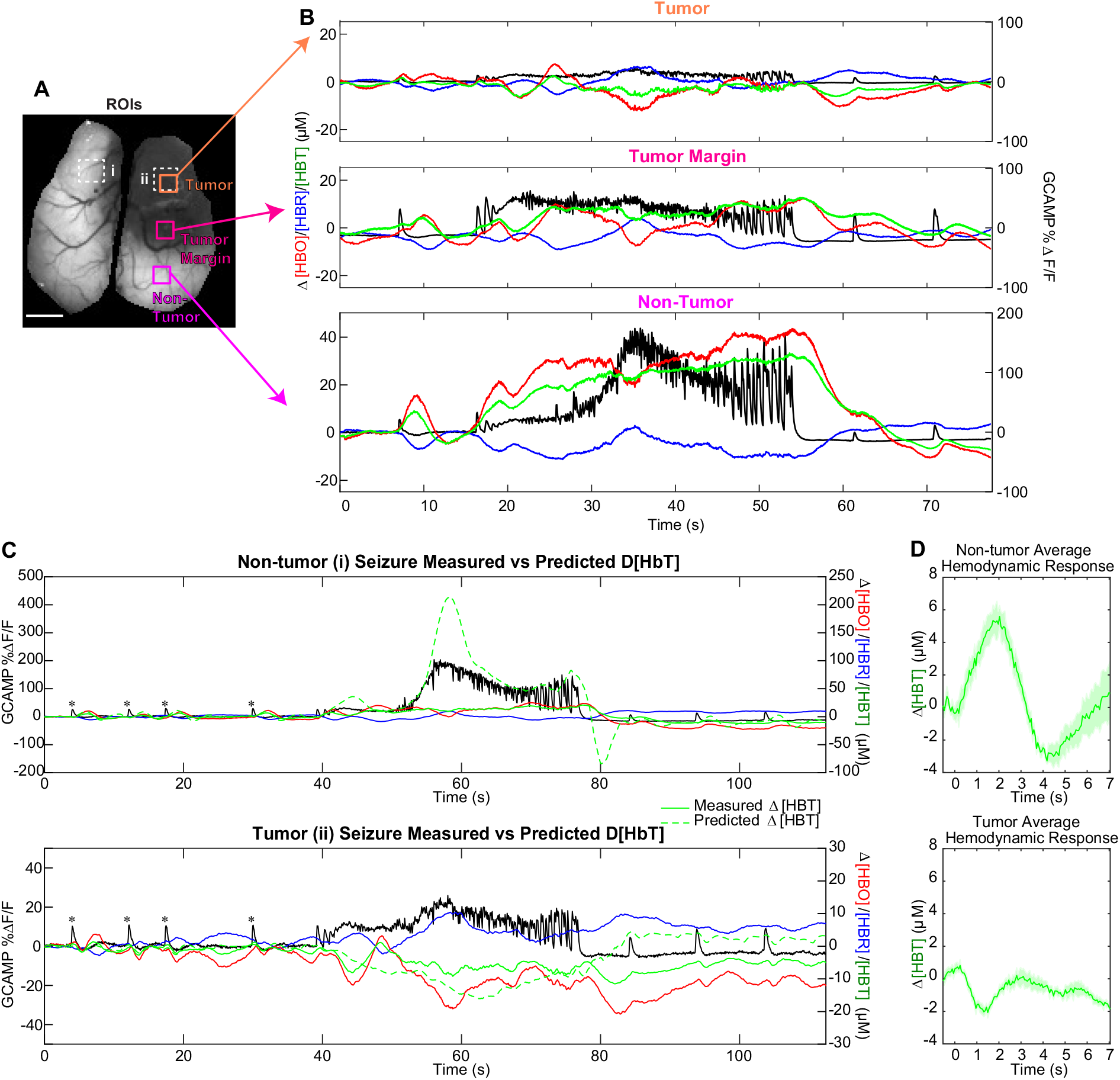
Seizures arise at the infiltrative margin and spread to tumor and non-tumor regions (Mouse 2). (A) ROIs over tumor region (orange), infiltrative margin (fuchsia) and non-tumor (magenta) areas used to obtain average time courses in (B) and over non-tumor and tumor region (light blue) for average time courses in (C). (B) Time series of generalized seizure event, shown for both GCaMP6f and hemodynamics in three regions; tumor (orange), infiltrative margin (fuchsia) and non-tumor regions (magenta). Vertical axes are scaled differently between GCaMP6f and hemodynamics to visualize all time-series but are consistent between tumor, margin and non-tumor time series displays. The seizure begins with multiple high-amplitude discharges at the infiltrative margin of the tumor and spreads to other cortical areas. (C) Time series of generalized seizure event, shown for both GCaMP6f and hemodynamics and averaged across non-tumor and tumor regions indicated as (i) and (ii) respectively in (A). Green solid lines represent measured change in total hemoglobin, and green dashed lines represent predicted change in total hemoglobin according to average hemodynamic response of four preceding interictal events (indicated with magenta asterisks) convolved with measured GCaMP6f time series. Vertical axes are scaled differently to visualize all time series. (D) Average hemodynamic response to four interictal events prior to seizure for non-tumor region (top) and tumor regions. Note that the non-tumor regions show a positive hemodynamic response whereas the tumor region shows a negative hemodynamic response. The standard deviation across four events is shown.

## Supplementary Videos

**Supplementary Video S1.**
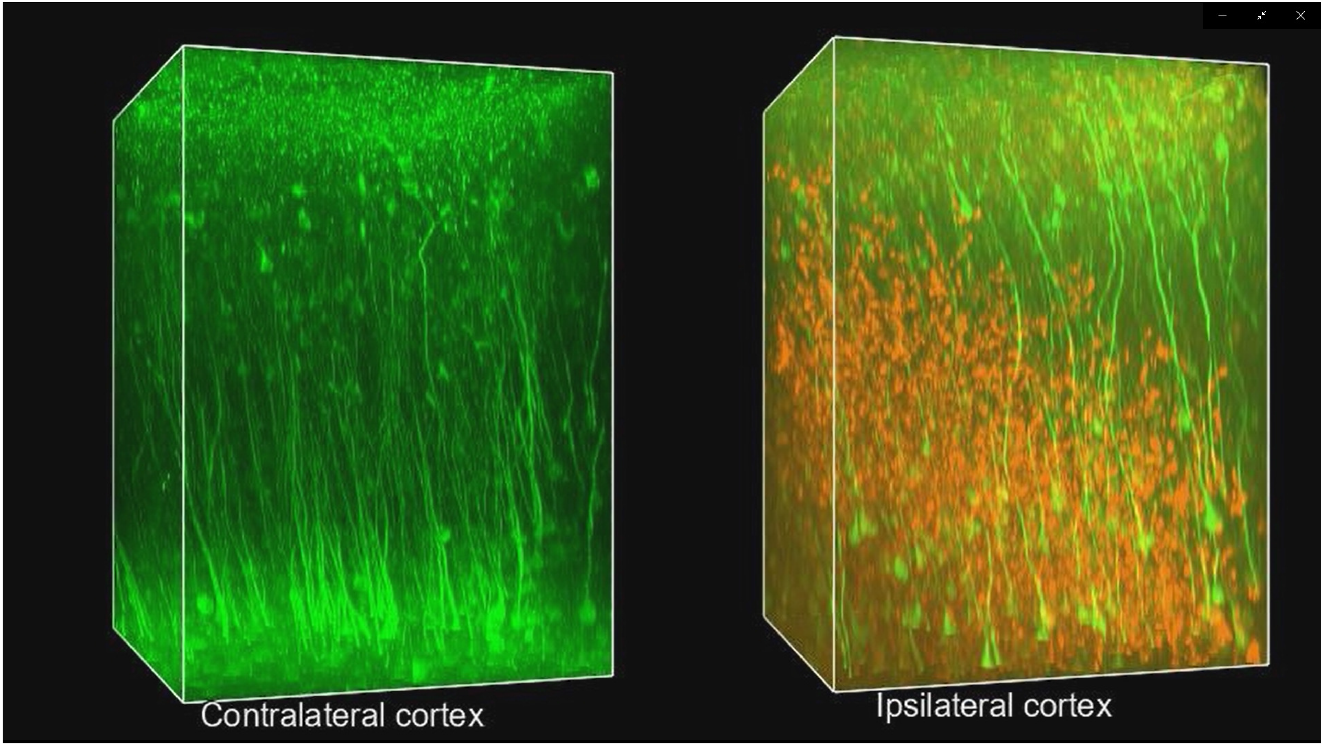
Histology comparison between tumor and contralateral cortex. Glioma cells intermingle with GCaMP6f+ neurons in the cortex. Video of three-dimensional renderings from the contralateral (left) and ipsilateral cortex (right) of mouse 1, acquired post-mortem using two-photon microscopy. Tumor cells (HA; red) and GCaMP6f+ neurons (green).

**Supplementary Video S2.**
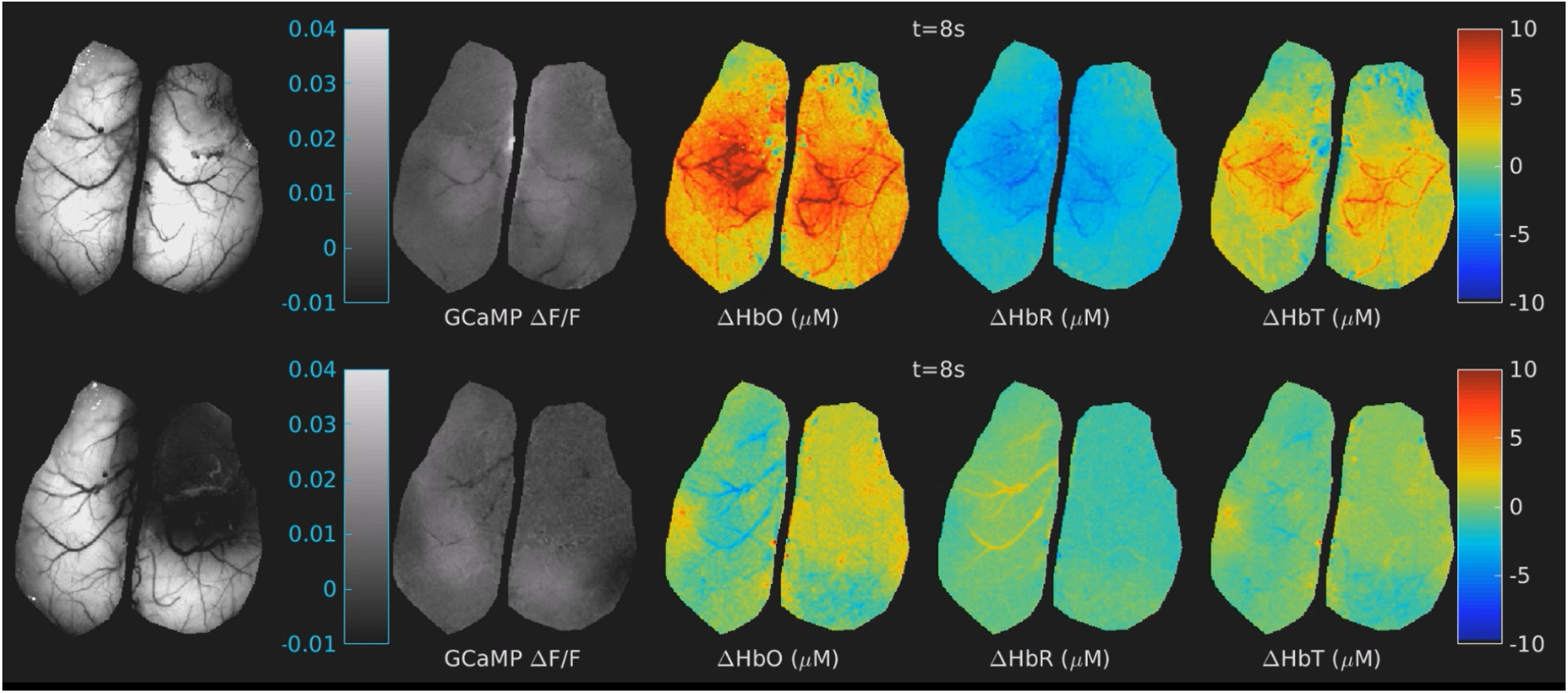
Resting-state neural and hemodynamic activity at 14 and 34 DPI. Examples of resting state neuronal and hemodynamic activity in earlier (14 days post injection; upper row) versus late (34 days post injection; bottom row) stage of glioma development in mouse 2. From left to right: raw fluorescence image, GCaMP6f fluorescence change, oxy-, deoxy- and total hemoglobin concentration change.

**Supplementary Video S3.**
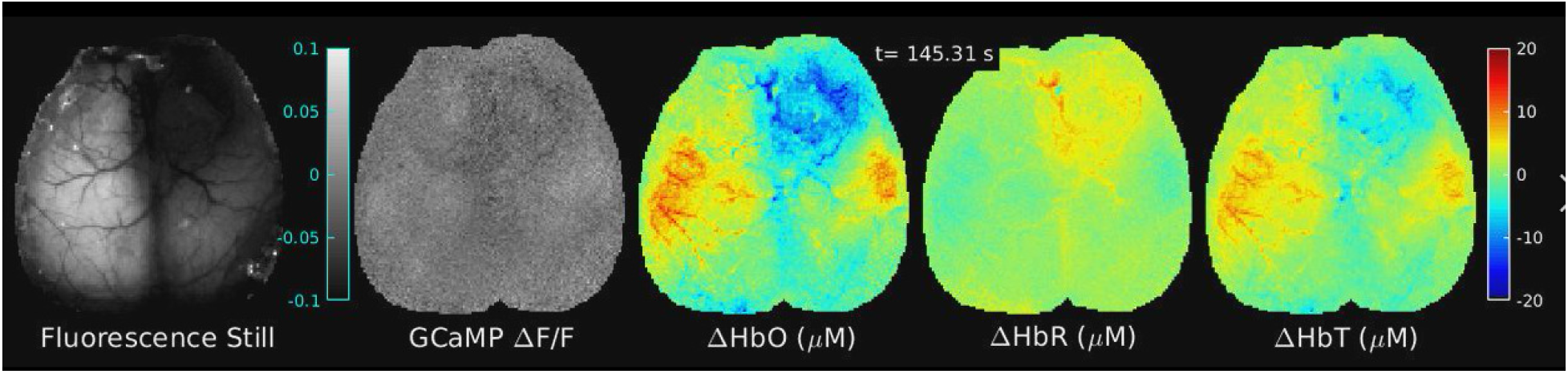
A full 180 second trial at 35 DPI showing 13 interictal discharges. 180 second full run from mouse 1 at DPI 35 showing 13 different inter-ictal events (same run as shown in Figure 4B-D). From left to right: Baseline fluorescence, GCaMP6f fluorescence change, oxy-, deoxy- and total hemoglobin concentration change. Note that the fixed color-scale compared to Video S2 is wider to allow for the high amplitude events, but smaller spontaneous fluctuations in neuronal activity and hemodynamics are still occurring.

**Supplementary Video S4.**
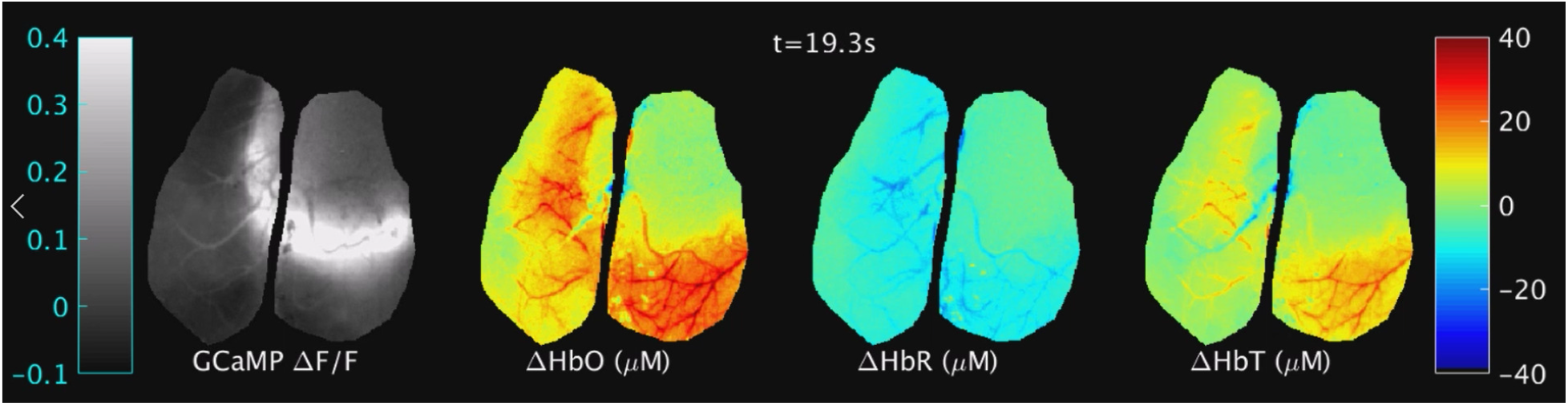
A real-time, spontaneous generalized seizure in mouse 1 at 36 DPI. Generalized seizure in mouse 1, observed 36 days post injection. Amplitudes of neuronal and hemodynamic activity are much lower (and for hemodynamics, sluggish and inverted) in the tumor area compared to very large non-physiological amplitudes in non-tumor areas. From left to right: GCaMP6f fluorescence change, oxy-, deoxy- and total hemoglobin concentration change.

**Supplementary Video S5.**
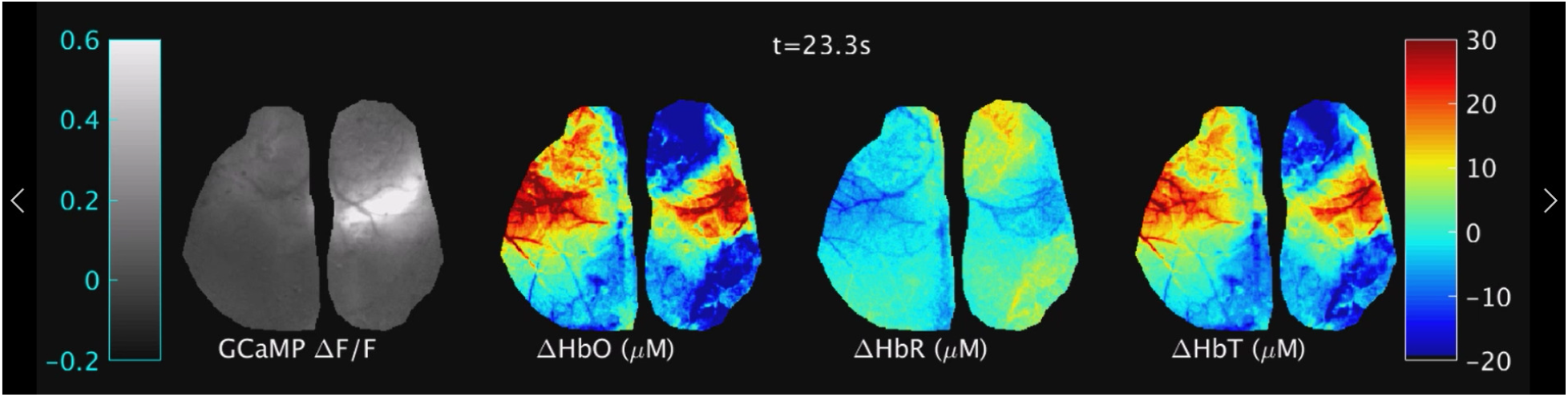
A real-time spontaneous generalized seizures in mouse 2 at 35 days DPI. Generalized seizure in mouse 2, observed 35 days post injection. Amplitudes of neuronal and hemodynamic activity are much lower (and for hemodynamics, sluggish and inverted) in the tumor area compared to very large non-physiological amplitudes in non-tumor areas. From left to right: GCaMP6f fluorescence change, oxy-, deoxy- and total hemoglobin concentration change.

